# Mapping Strigolactone Hydrolysis in DWARF14 via QM/MM String Method

**DOI:** 10.1101/2022.04.22.489200

**Authors:** Tanner J. Dean, Jiming Chen, Diwakar Shukla

**Affiliations:** Center for Biophysics and Quantitative Biology, University of Illinois at Urbana-Champaign, Urbana, IL 61801; Department of Chemical and Biomolecular Engineering, University of Illinois at Urbana-Champaign, Urbana, IL 61801; Department of Bioengineering, University of Illinois at Urbana-Champaign, Urbana, IL 61801; Department of Plant Biology, University of Illinois at Urbana-Champaign, Urbana, IL 61801; Department of Chemistry, University of Illinois at Urbana-Champaign, Urbana, IL 61801

## Abstract

Strigolactone signaling is unusual among plant hormone pathways in that its receptor, DWARF14 (D14), also catalyzes the hydrolysis of the hormone. However, key aspects of this reaction remain unresolved including (i) whether hydrolysis proceeds via the canonical acyl substitution method initiated by nucleophilic attack on the butenolide (D-ring) or an alternative Michael addition at the enol-ether bridge, and (ii) the identity of the hydrolysis-induced covalent modification that promotes receptor activation. In this work we test these competing mechanistic hypotheses using QM/MM string method simulations of the D14-GR24 complex. Our simulations show that the acyl substitution pathway initiated by nucleophilic attack of the D-ring is strongly favored, supporting the canonical mode proposed by experimental studies. Furthermore, we find that multiple covalent adducts involving the butenolide ring and catalytic residues can form and interconvert along the reaction coordinate. These results suggest that the hydrolysis-induced covalent modification is not a single static species, but a dynamic ensemble of chemically related states. Our findings provide a mechanistic framework that reconciles previously conflicting experimental observations and clarifies the chemical basis by which strigolactone hydrolysis promotes receptor activation.

## Introduction

Strigolactones are a class of plant hormones responsible for regulating shoot branching, root architecture, and hypocotyl elongation in plants.^1–6^ In flowering plants, they are perceived a class of receptors known as DWARF14 (D14) proteins.^7^ A structure of a D14 protein is shown in Fig. 1A. Strigolactone signaling begins with binding of the ligand to D14 followed by catalytic hydrolysis (Fig. 1D, steps 1-2).^8,9^ After the strigolactone molecule is hydrolyzed a portion remains covalently bound to the catalytic residues while the receptor undergoes a large conformational change from its “open” or inactive state to its “closed” or active state (Fig. 1D, step 3).^10,11^ While in its active state, the receptor associates with its signaling partner MAX2 as well as a SMXL protein (Fig. 1D, step 4).^12,13^ The complex is then ubiquitinated and degraded, inducing further downstream signaling (Fig. 1D, steps 5-6).^14^ One component of strigolactone perception by D14 proteins is hydrolysis of the substrate by D14. While the strict dependence of downstream signaling on substrate hydrolysis has been disputed,^15^ this hydrolysis reaction has been shown to be a promoter of signaling. ^10,16,17^ Intriguingly, while a strigolactone receptor in parasitic witchweed, *Sh*HTL7, has been shown to have a unique picomolar EC50 for signaling in the presence of strigolactone analog GR24,^18^ its *apo* conformational dynamics suggest it has lower signaling activity compared to D14. ^19^ This further suggests that substrate hydrolysis and its resulting covalent modification of the receptor plays a role in activating these receptors and highlights the importance of understanding the details of the strigolactone hydrolysis mechanism.

**Figure 1:**
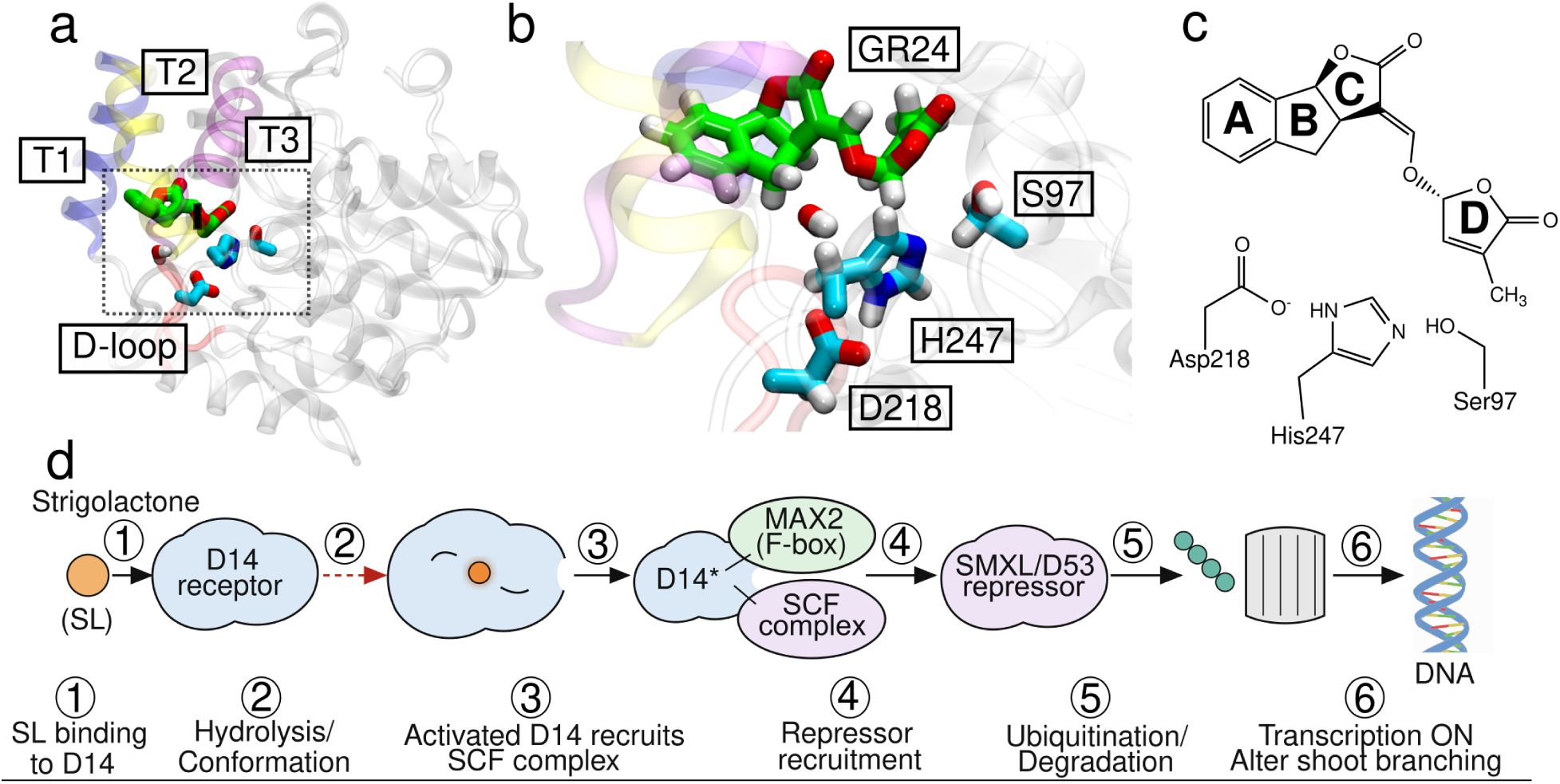
(A) Overall structure of *At* D14 with the T1, T2, and T3 helices shown in blue, yellow, and purple respectively and D-loop shown in red. The catalytic triad, GR24 ligand, and a binding pocket water molecule. The boxed region contains the QM atoms. (B) The QM region consisting of S97-D218-H247 catalytic triad, GR24 (strigolactone analog) ligand, and binding pocket water. (C) Schematic of GR24 and the catalytic triad. GR24 is shown in its 5DS stereochemistry, matching that of naturally occurring strigolactones. (D) The strigolactone plant hormone pathway. While steps 1 and 3-6 are well studied, the exact mechanism of catalysis in step 2 remains unknown.

The canonical model for strigolactone hydrolysis is that hydrolysis begins via a nucleophilic attack of the catalytic S97 residue on the butenolide ring (D-ring) of strigolactone.^10^ However, it has also been suggested that the reaction may occur via a Michael addition pathway in which the nucleophilic attack instead occurs on the enol-ether bridge.^20^ Reaction schemes for these proposed pathways are shown in Fig. 2. In addition, while several studies have indicated that the D-ring is covalently bound to the protein when it undergoes its conformational transition to the active state, there is disagreement on the form and location of the D-ring.^10,17,21,22^ Mass spectrometry results of a peptide containing residues surrounding H247 from RMS3, a D14 ortholog, showed a species matching the mass of butenolide ring bound to H247.^17^ A crystal structure of the active form of the protein showed what was believed to be a covalently linked intermediate molecule (CLIM), which is an open butenolide ring covalently bound to both S97 and H247 of the catalytic triad.^10^ Reanalyses of this crystal structure has shown that the electron density attributed to CLIM is also able to fit well with an iodide ion^23^ or a closed D-ring bound to the catalytic histidine (D-ring-H247).^21^

**Figure 2:**
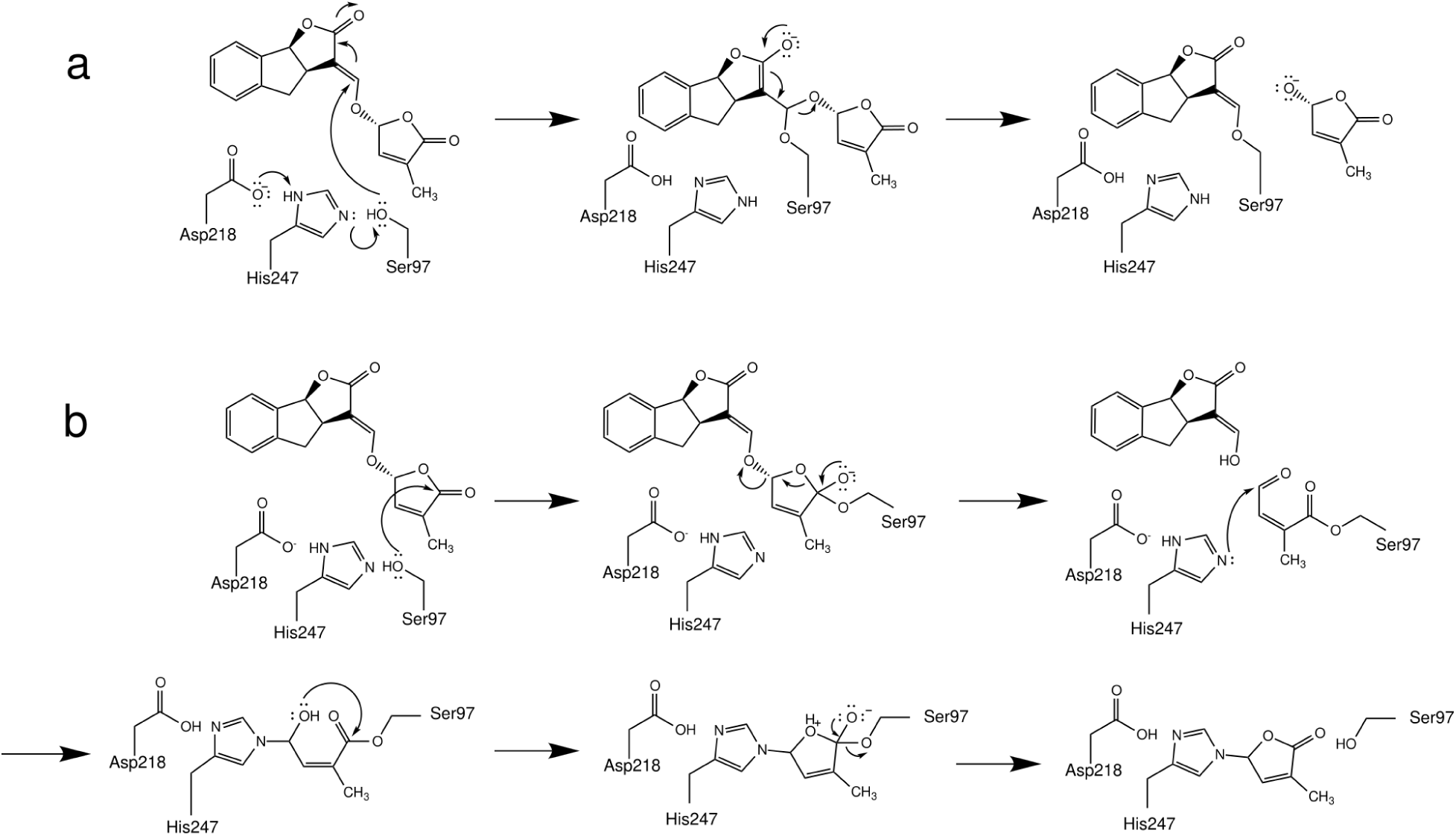
Reaction pathways for the (A) Michael addition and (B) acyl substitution reaction pathways proposed in literature. The Michael addition pathway is initiated by a nucleophilic attack on the enol-ether bridge of GR24, while the acyl substitution pathway is initiated by a nucleophilic attack on the carbonyl carbon on the butenolide ring (D-ring) of GR24. This pathway includes covalently linked intermediate (CLIM) species and covalent D-ring H247 modification (D-ring-H247) that have been reported in literature.

Taken together, these studies highlight two central unresolved questions regarding the mechanism of strigolactone hydrolysis:

1. What is the dominant reaction pathway for strigolactone hydrolysis?

(1A) Does hydrolysis proceed via the canonical acyl substitution mechanism initiated by nucleophilic attack on the butenolide ring (D-ring)?
(1B) Does hydrolysis proceed via the alternatively proposed Michael addition pathway involving attack at the enol-ether bridge?
2. What is the nature of the hydrolysis-induced covalent modification that promotes receptor activation?

1. (2A) Is there a single dominant covalent species (e.g., CLIM or D-ring-H247)?
2. (2B)Does hydrolysis give rise to multiple interconverting covalent intermediates?

In our previous work, we have used long-timescale molecular dynamics simulations to characterize substrate recognition and conformational dynamics of strigolactone receptors, ^19,24^ studied how the *Os*D14-*Os*D3 C-terminal helix complex formation effects binding of strigolactones,^25^ and predict the effects of histidine modification on active state stability in *At* D14 and *Sh*HTL7.^26^ However, a limitation to classical molecular dynamics is that they are unable to capture changes in covalent bonds such as those involved in chemical reactions. Thus, to address the questions outlined above, we employed QM/MM free energy simulations to characterize the proposed reaction pathways in detail. Briefly, QM/MM is a molecular simulation method in which a portion of a system is described using quantum mechanical calculations, while the rest of the system is described using a classical molecular dynamics force field.^27–29^ This enables simulations of enzymatic systems that would be intractable by pure quantum mechanical calculations. A schematic of our quantum mechanical region is shown in Fig. 1B-C. Since QM/MM simulations have a high computational cost, we employed string method free energy calculations to determine optimal reaction pathways for each mechanistic step. The string method is a transition path search method a path is discretized into images and the images are updated iteratively using restrained simulations.^30–32^ This method has been previously used in conjunction with QM/MM simulations to study other enzymatic systems.^33–36^ String method optimization additionally enables computation of free energy profiles along reaction pathways, which in turn enables identification of the most probable of a set of possible reaction pathways. In this manuscript, we employed extensive string method free energy calculations (7 string optimizations with ∼1.7 ns aggregate simulation time at DFT/B3LYP-level theory) on previously proposed mechanisms of strigolactone hydrolysis. Using our simulation results, we have been able to identify the favored reaction mechanism and reconcile partially conflicting models of hydrolysis-induced covalent modification of strigolactone receptors.

## Results

### Acyl substitution pathway is energetically favored

Two possible mechanisms of strigolactone hydrolysis have been proposed in the literature: a Michael addition pathway in which the S97 attacks the enol-ether bridge, ^20^ and an acyl substitution pathway in which S97 attacks the butenolide ring.^10^ To determine the more probable reaction pathway for strigolactone hydrolysis, we performed QM/MM string optimizations on each reaction step of both pathways. In reaction pathways, the step with the highest free energy barrier was the initial nucleophilic attack of S97 on the substrate. This indicates that the rate-limiting process for GR24 hydrolysis is the S97 attack. The free energy barrier of the initial S97 attack on the substrate is ∼14 kcal/mol in the acyl substitution pathway, while the barrier for the initial attack is ∼16 kcal/mol in the Michael addition mechanism (Fig. 3-4). This nucleophilic attack is highly endothermic in both pathways with Δ*G* values of ∼11.3 kcal/mol and ∼12.7 kcal/mol in the Michael addition and acyl substitution pathways, respectively. These free energy profiles indicate that the acyl substitution pathway is kinetically more favorable than the Michael addition pathway and very similar thermodynamically. Error bars for these PMF profiles are shown in Tables SS4-S10.

**Figure 3:**
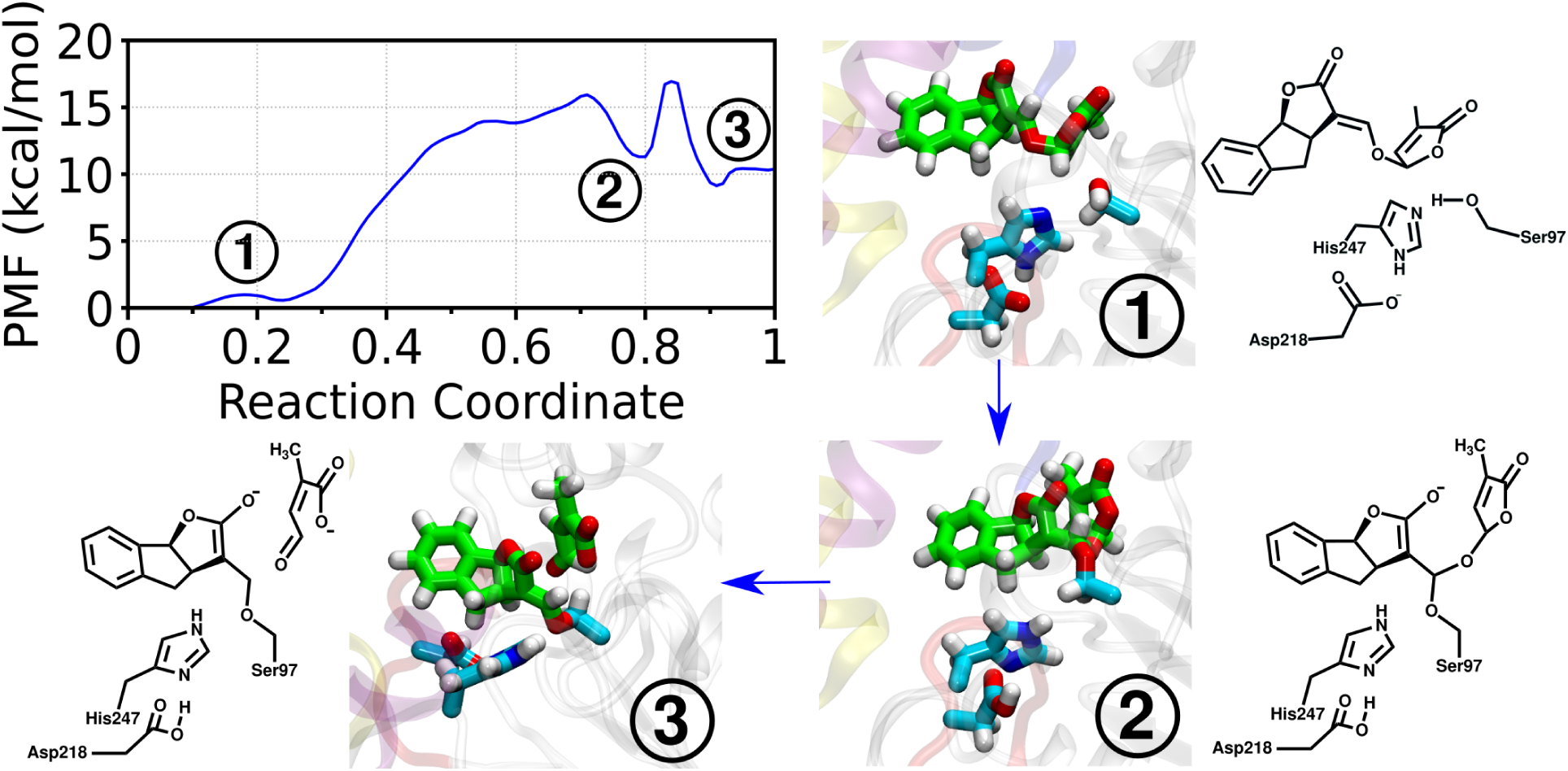
Potential of mean force profile for the Michael addition pathway calculated from simulations. The structures of (1) the initial GR24 and catalytic triad, (2) intermediate species with GR24 covalently bound to S97 via the enol-ether bridge, and (3) the final state consisting of the ABC-rings and detached, open-form D-ring are shown, and their locations on the PMF profile are indicated with their corresponding numbers.

**Figure 4:**
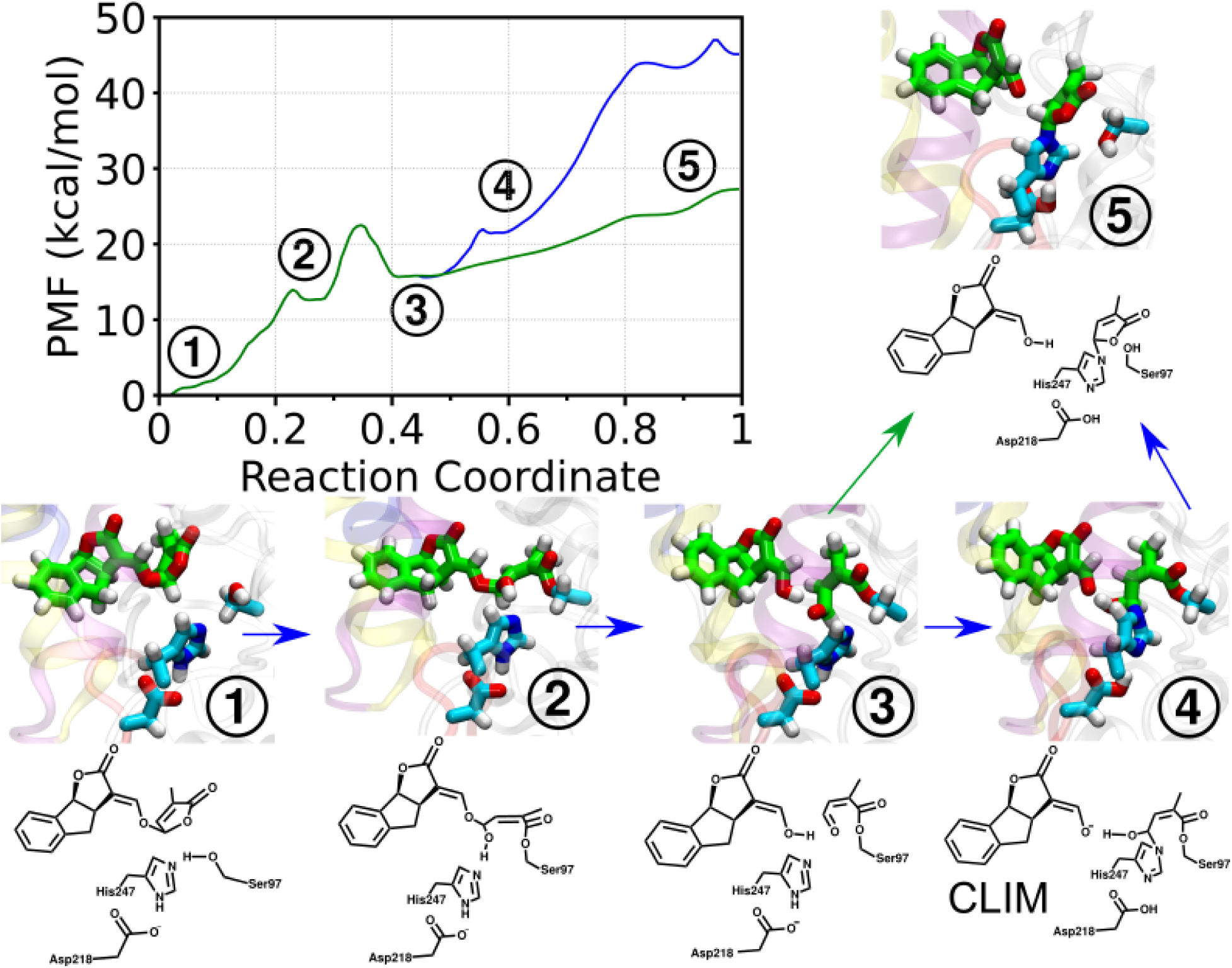
Potential of mean force profile for the acyl substitution pathway calculated from simulations. The structures of (1) the initial GR24 and catalytic triad, (2) intermediate with GR24 covalently bound to S97 via the D-ring, (3) intermediate with the open D-ring covalently bound to S97, (4) covalently linked intermediate (CLIM), (5) and closed D-ring covalently bound to H247 are shown, and their locations on the PMF profile are indicated with their corresponding numbers.

Following the initial nucleophilic attack, the enol-ether bridge is severed so that the butenolide ring is detached from the ABC ring scaffold. In the Michael addition pathway, the ABC ring scaffold remains covalently bound to the serine after hydrolysis, while the D-ring is not covalently bound to any of the catalytic triad residues. In contrast, the D-ring remains covalently bound to either S97 or H247 post-hydrolysis. Several studies have detected D-ring derivatives that remain covalently bound to the enzyme, which is consistent with our finding that the acyl substitution pathway is more favorable. Additionally, fluorescent strigolactone analogs which are often used to study strigolactone receptors *in vitro* contain butenolide rings but lack an enol-ether bridge, suggesting that an enol-ether bridge is not needed for the substrate to be hydrolyzed.^37^ This further suggests that strigolactone hydrolysis follows the acyl substitution pathway. Mechanistic details of the acyl substitution pathway are discussed in the following section, and details of the unfavored pathways, Michael addition and the D-ring-H247 formation from CLIM step from the acyl substitution pathway, are shown in Figs. S1-S3. Observed mechanisms with arrow-pushing schematics are shown in Figs. S4-S10.

### Mechanistic details of acyl substitution

#### Nucleophilic attack on D-ring leads to spontaneous ring opening

The acyl substitution pathway begins with a nucleophilic attack of S97 on the C5’ carbon of GR24 (Fig. 5A). This nucleophilic attack is facilitated by a proton transfer from S97 to the adjoining H247 of the catalytic triad, which increases the nucleophilic strength of the serine. This reaction step is endothermic by ∼12.7 kcal/mol with a free energy barrier of ∼14 kcal/mol. During the initial steered MD and string optimization of this reaction step, biases were applied to the proton transfer and the nucleophilic attack distance. In addition to these bonds, two additional bond rearrangements occurred spontaneously, without applied biases: proton transfer from H247 to an oxygen on the D-ring of GR24 and opening of the D-ring of GR24. The occurrence of this ring opening process without the application of applied forces suggests that the free energy barrier associated with the ring opening is insignificant compared to other reaction barriers. Fig. 5B shows distances between key bonds that form and break for each string image for this step. These distances show that formation of the S97-GR24 bond coincides with transfer of a proton between S97 and H247. The opening of the D-ring also coincides with a transfer of the same proton from the H247 to an oxygen on the open D-ring. In a study of strigolactone analogs, carba-SL compounds with modified butenolide rings that prevent separation of the D-ring from the ABC-rings were tested as strigolactone antagonists. A crystal structure of D14 in complex with one such compound found the unhyrolyzed compound with an open-form D-ring covalently bound to S97.^38^ This crystollographic evidence is consistent with our finding that the D-ring can open upon nucleophilic attack by the serine prior to separation of the D-ring from the ABC rings.

**Figure 5:**
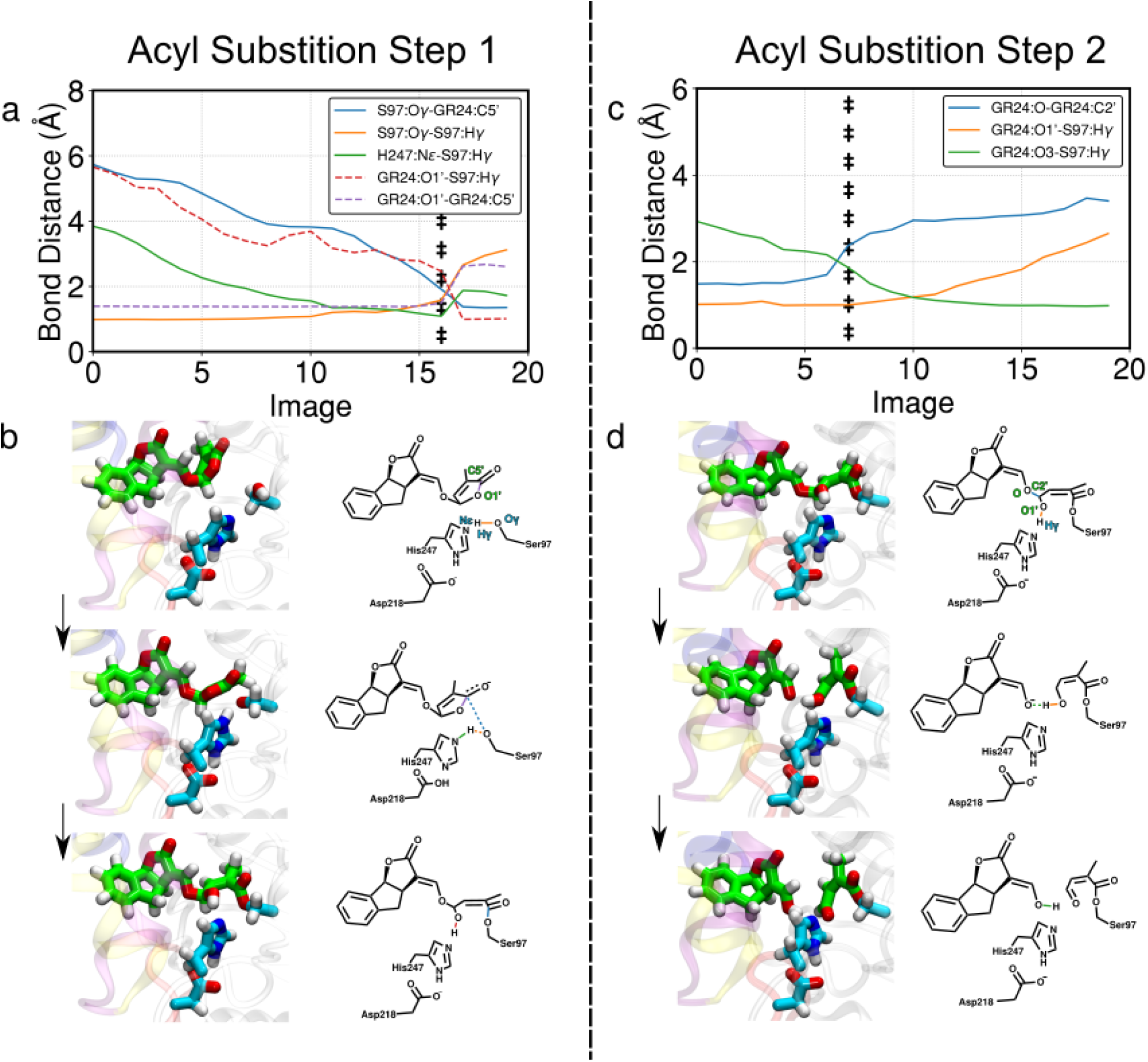
Distances between atoms that form and break bonds during the initial nucleophilic attack of Ser97 on the D-ring and the ABC-ring separation from the D-ring. (A,C) Solid lines indicate distances that were constrained during string optimization, and dashed lines indicate bonds that changed spontaneously without applied biases. The transition state, defined as the maximum point on the PMF for the step, is demarcated by double daggers. s(B) Structures of the (left to right) initial state of the intact GR24 substrate with the catalytic triad transition state, and final state with GR24 covalently bound to S97 are shown. (D) Structures of the initial state with GR24 covalently bound to S97, transition state, and final state with an open D-ring covalently bound to S97 are shown.

#### ABC separation leads to modification of S97 with D-ring

After the S97 residue forms a bond with the C5’ carbon on the D-ring, the enol-ether bridge of GR24 is severed. This occurs via conversion of the hydroxyl group on the C12 carbon to a carbonyl and a coupled proton transfer (Fig. 5)B. This step is ∼3 kcal/mol endothermic with a barrier of ∼10 kcal/mol. The resulting intermediate species is an open D-ring on S97 (Open D-S97), which is hypothesized to be the precursor to a covalently linked intermediate (CLIM) molecule proposed by Yao *et al*.^10^ While our simulations were of insufficient length to capture the ABC rings unbinding, the ABC rings could, in principle, unbind from the enzyme active site at this stage since it is not covalently bound any of the catalytic triad residues or the D-ring. Previous work has additionally found that the ABC rings are partially solvent-exposed in resolved crystal structures, consistent with it being a leaving group. ^12^

### Multiple possible modes of covalent modification to H247

Previous studies have suggested that covalent modification to H247 of the catalytic triad acts as a promoter of signaling activity. However, there is disagreement over whether this modification is the CLIM, which is an open D-ring linked to both S97 and H247, or a closed butenolide ring attached only to H247 (D-ring-H247). To evaluate these possibilities, we ran string optimizations from Open D-S97 to CLIM to D-ring-H247 as well as Open D-S97 to D-ring-H247. We find that both CLIM and D-ring-H247 can be formed upon nucleophilic attack of H247 on the carbonyl group of Open D-S97.

Fig. 6A shows the reaction to form a CLIM from Open D-S97. This involves H247 performing a nucleophilic attack on the C2’ carbon on the open D-ring and is coupled with protonation of the O1’ oxygen of the D-ring. In our simulations, the source of this proton was the hydroxyl group attached to the ABC rings, however, in the absence of the ABC rings, water could act as a proton source as well. Formation of the CLIM is a ∼5 kcal/mol process with a free energy barrier of ∼6.2 kcal/mol. The reaction mechanism proposed by Yao *et al.* predicts that after CLIM formation, the O1’ oxygen on the D-ring performs a nucleophilic attack on the C5’ carbon, which induces the dissociation of the D-ring from S97. However, our string optimization of the conversion from CLIM to D-ring-H247 suggests that the D-ring first dissociates from S97 prior to ring closure (Fig. S3). This has an energy barrier of ∼25.5 kcal/mol, which suggests that the CLIM is highly stable.

**Figure 6:**
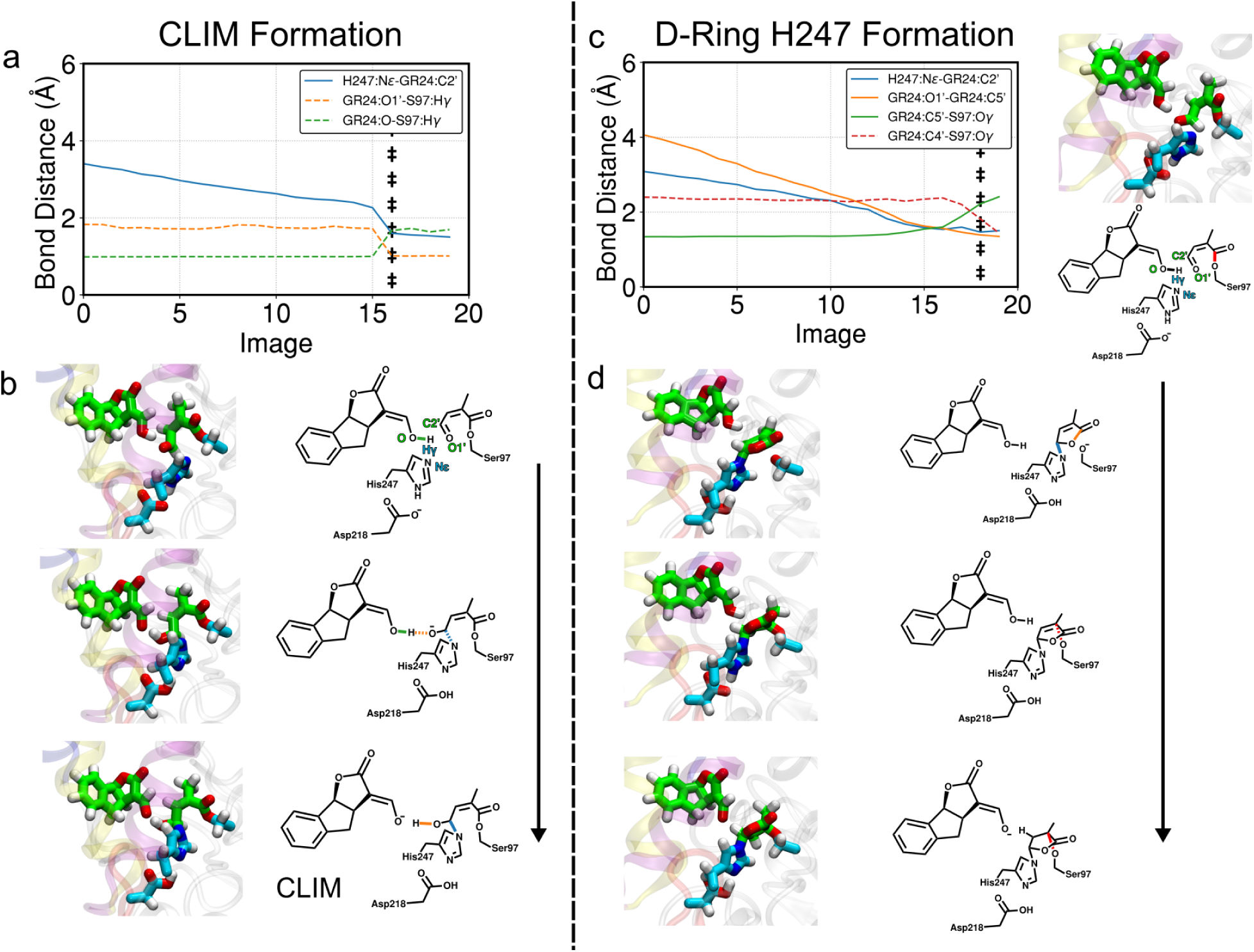
Distances between atoms that form and break bonds during covalently linked intermediate (CLIM) or D-ring-H247 formation. (A,C) Solid lines indicate distances that were constrained during string optimization, and dashed lines indicate bonds that changed spontaneously without applied biases. The transition state, defined as the maximum point on the PMF for this step, is demarcated by double daggers. (B) Structures of the initial state with the open D-ring covalently bound to S97 (Open D-S97), transition state, and CLIM species bound to both S97 and H247 are shown. (D) Structures of the initial state with the open D-ring covalently bound to S97 (Open D-S97), D-ring-H247 species observed in image 16 of the string, transition state, and closed form of D-ring bound to both S97 and H247 are shown.

Alternatively, we evaluated the possibility of a concerted formation of D-ring-H247 from Open D-S97 without forming a stable CLIM intermediate. In this pathway, we observed that formation of D-ring-H247 is a monotonically uphill process with an energy barrier of ∼11.5 kcal/mol. Following formation of this species, S97 can form a covalent bond with the C4’ carbon of the D-ring. The activation barrier to forming this bond is low (∼1 kcal/mol) and the resulting species is very close in relative free energy to D-ring-H247 (*<*1 kcal/mol difference). Due to both this low free energy barrier and small difference in relative free energies, these species are likely to interconvert rapidly until S97 is protonated and thus weakened as a nucleophile. While there were no water molecules near S97 in our simulation system due to the presence of the ABC rings, the binding pocket is likely to fill with water when the ABC rings unbind, as indicated by several crystal structures of D14 homologs showing water in the binding pocket.^39–41^ This would enable S97 to be protonated, shifting the equilibrium toward the D-ring-H247 species. Detailed analysis of this pathway indicates that there is a transient CLIM-like species which forms (Fig. S9), however, the ring quickly closes in a positive ∼3 kcal/mol process in which the O1’ oxygen of the D-ring attacks the C5’ carbon, forming a tetrahedral intermediate. According to Baldwin’s rules for ring closure reactions,^42,43^ reformation of the five-membered butenolide ring is a favored 5-exo-trig reaction. In this nomenclature, “5” denotes formation of a five-membered ring, “exo” indicates that the nucleophilic attack occurs from outside the forming ring framework, and “trig” refers to an attack at a trigonal (sp^2^-hybridized) carbon center. In the present case, the O1’ oxygen attacks the sp^2^-hybridized C5’ carbon from an exocyclyic position, which is geometrically favorable and leads to rapid ring closure, forming a tetrahedral intermediate at C5’. The tetrahedral center then reconverts to a trigonal center by conversion of an oxygen to a carbonyl group, which induces dissociation of the D-ring from S97. The remaining species is a closed butenolide ring covalently bound to H247.

Taken together, these results suggest that while CLIM formation is chemically feasible, it is unlikely to represent an obligate intermediate along the productive pathway to the D-ring-H247 product. Although CLIM formation from Open D-S97 proceeds with a modest free energy barrier, the subsequent conversion of CLIM to D-ring-H247 is associated with a substantially higher barrier (∼25.5 kcal/mol), rendering CLIM a deep local minimum on the free energy surface. In contrast, direct formation of D-ring-H247 from Open D-S97 occurs through a lower-barrier, monotonically uphill pathway and is therefore kinetically favored. These observations support a model in which CLIM represents an off-pathway, kinetically trapped intermediate.

## Discussion

Using QM/MM free energy simulations, we characterized proposed mechanisms of strigolactone hydrolysis. Based on the free energy barriers of the initial nucleophilic attack of S97 upon the GR24 molecule, we have determined that the reaction occurs via an acyl substitution pathway rather than a Michael addition pathway. This is corroborated by experimental detection of intermediates associated with the acyl substitution pathway via mass spectrometry^17^ and X-ray crystallography experiments.^10^ Additionally, strigolactone analogs have previously been developed as synthetic agonists and antagonists for strigolactone receptors.^38,44,45^ In developing these analogs, direct crystallographic evidence of S97 forming a bond with the butenolide ring has been observed.^38^ It has also been observed that presence of a butenolide ring enhances binding and bioactivity.^44,46^ These suggest that the butenolide ring is a key portion of the substrate recognition and hydrolysis processs. In contrast, some strigolactone analogs such as Yoshimulactone Green, a fluorenscent analog commonly used to study strigolactone hydrolysis *in vitro*, undergoes hydrolysis despite it lacking an enol-ether bridge.^37^ This further suggests that hydrolysis occurs via nucleophilic attack on the butenolide ring rather than the enol-ether bridge.

In addition, we addressed the disagreement in the field regarding whether the covalent modification to the receptor that promotes receptor activation is the CLIM that is bound to both S97 and H247 or a closed butenolide ring that is only bound to H247 (D-ring-H247). Our results indicate that while formation of either species is possible, the free energy barrier to forming a CLIM is lower, indicating that a stable CLIM is likely the dominant covalent modification present. However, the D-ring-H247 modification is able to form either directly from an Open D-S97 species that forms after D-ring separation from the ABC rings, or from a CLIM, though the high free energy barrier of the latter suggests that it is highly improbable. Formation of the D-ring-H247 modification occurs via a 5-exo-trig ring closure, which is favored per Baldwin’s rules.^42,43^ During this conversion, a CLIM-like species is observed (Fig. S9), however, this species differs from the stable CLIM in several key ways. The first is that the O1’ oxygen atom that performs an intramolecular nucleophilic attack to close the D-ring is deprotonated while bonded directly to an sp3-hybridized carbon, thus giving it a negative charge. Since negative charge is expected to enhance nucleophilic strength, ^47,48^ the O1’ oxygen in this CLIM-like species is expected to have greater nucleophilic strength than in the stable CLIM, where the O1’ oxygen is protonated. The other key difference is proximity of the O1’ oxygen to the C5’ carbon it nucleophilically attacks to close the D-ring. From Fig. 6C, the distance between the O1’ and C5’ atoms of GR24 are ∼2 Å apart at the image where the CLIM-like state is observed (image 14), whereas in the stable CLIM they are ∼4 Å apart. It has been suggested that reaction rate in intramolecular reactions is highly sensitive to distance between reactive species, with changes in distance less than 1 Å sometimes leading to orders-of-magnitude changes in reaction rates.^49,50^ Thus, it is likely that closer proximity of the O1’ and C5’ in the CLIM-like species compared to the stable CLIM is a strong contributing factor to its propensity to perform a ring closure reaction and form the D-ring-H247 species. Our simulations were also able to capture D-ring-H247 formation from the stable CLIM, however, the reaction happens via a different pathway than shown in Fig. 2B. Rather than a direct 5-exo-trig ring closure as seen in the pathway from Open D-S97 to D-ring-H247, the D-ring first dissociates from S97 and forms a covalent bond via a different carbon atom (Fig. S10), which is a precursor to a 5-endo-dig ring closure that closes the D-ring. While this is considered to be a favorable ring closure per Baldwin’s rules and known to form butenolide rings,^51^ the PMF profile for the formation of D-ring-H247 from CLIM indicates that there is a ∼24 kcal/mol barrier to forming the precursor. Thus, the D-ring-H247 modification is more likely to occur via a concerted mechanism from the Open D-S97 species since the free energy barrier is lower, at ∼11.5 kcal/mol.

Previous literature on the nature of the covalent modification include mass spectrometry showing a D-ring bound to H247,^17^ a crystal structure of a CLIM species,^10^ reanalysis of the CLIM structure suggesting that the electron density fits will with an intact D-ring,^21^ and a crystal structure showing a D-ring on the catalytic serine in KAI2, a homolog of D14 with the same catalytic triad.^52^ While these all initially appear to be competing views, our simulations show that all of these covalent modifications are likely to be present in some quantity. Both the CLIM and D-ring-H247 modifications are able to form during substrate hydrolysis. Additionally, the pathway to forming the D-ring-H247 contains a transient CLIM-like species, as previously discussed, and a transient closed D-ring bound to both S97 and H247. After the D-ring dissociates from S97 while forming the D-ring-H247 species, S97 can reattach to the D-ring while still deprotonated (Fig. S9). In principle, any of these interconverting covalent modifications could contribute electron density in a crystal structure, thus it is plausible that different forms of the D-ring covalent modification on the S97 and H247 residues are detected in different crystal structures. Additionally, since the D-ring is able to dissociate from and reassociate with S97 (Fig. S9), mass spectrometry on a peptide containing H247 but not S97, as performed by de Saint Germain *et al.* would be expected to show the presence of a butenolide ring. A constitutively active D218A mutant of D14 in which the D218-H247 interaction in the catalytic triad is disrupted^15^ suggests that modification to H247 would act as a promoter of strigolactone receptor activation. Thus, any of the present forms of the D-ring, so long as they are covalently bound to H247, are likely to act as promoters of strigolactone receptor activation as well.

Taken together, this work provides a unified mechanistic framework for strigolactone hydrolysis that resolves several long-standing ambiguities in the field. By direclty comparing competing pathways usign QM/MM free energy simulations, we establish that hydrolysis proceeds via the canonical acyl substitution mechanism, thereby clarifying the chemical origin of catalysis in D14 receptors. In addition, our results redefine the role of the covalently-linked intermediate, showing that while it is chemically accessible, it does not constitute an obligate intermediate along the dominant reaction pathway and instead represents a kinetically trapped off-pathway state. More broadly, our findings support a model in which hydrolysis generates a dynamic ensemble of interconverting covalent modifications rather than a single static species. This perspective reconciles previously conflicting structural and mass spectrometry observations and suggests that receptor activation is driven by the presence of the D-ring modification on H247, independent of its precise chemical form. By resolving the reaction mechanism and the nature of the hydrolysis-induced modification, this study provides a foundation for the rational design of strigolactone analogs that modulate receptor activity.

## Methods

### System preparation

The structure of *At* D14 was obtained from PDB 4IH4.^53^ The GR24 ligand was superimposed into the binding pocket using the GR24-bound structure of *Os*D14 as a template (PDB code 5DJ5)^12^ in PyMOL.^54^ Parameters for GR24 were produced using the CHARMM General Force Field (CGenFF).^55^ The complex was solvated in a TIP3P water box with the VMD solvate plugin and a 0.15 M NaCl concentration was added with the VMD plugin autoionize. Following initial preparation of parameters for the separate system sections, VMD was used to produce the psf and structure files necessary for the simulations.^56^ The quantum mechanical region was defined as the GR24 ligand, the catalytic triad, and water molecules in the binding pocket and described using the B3LYP functional^57^ and 6-31G* basis set^58^ with a separate psf and structure file. The molecular mechanical (MM) region was described using the CHARMM36 force field.^59^

### Simulation protocol

Prior to QM/MM simulation, the system was minimized using the conjugate gradient descent method for 10000 steps classically, followed by a 5 ns classical equilibration to allow water molecules to equilibrate. After this classical equilibration, the system was minimized using a QM/MM scheme for 1000 time steps again using conjugate gradient descent and equilibrated for 1 ps. Production runs were run at 300 K and 1.0 bar using a Langevin thermostat and Langevin barostat.^60,61^ Following initial equilibration, initial reaction paths for each proposed mechanism were generated using steered molecular dynamics simulations using bonds formed and broken at each proposed mechanistic step as reaction coordinates. ^62^ A full table of reaction coordinates is shown in Table S1. Following each steered MD simulation, a 1 ps equilibration was performed before generating the steered MD pathway for the subsequent mechanistic step. Steered MD trajectories were used for string optimization. Classical equilibration simulations were performed using NAMD 2.13, ^63,64^ and QM/MM simulations were performed using the NAMD QM/MM interface^65^ in conjunction with TeraChem.^66^

### String optimization

For string optimization of each mechanistic step, each pathway was discretized into 20 im-ages. Images were restrained on the same collective variables used for steered MD. Restrained simulations were performed for each image for ∼100 fs with 5 replicates per image. Details of the simulation time for each string optimization are shown in Tables S2-S3. Following each step, the restraint centers for each image were updated as follows: (i) Drifted centers were computed using the average values of collective variables for each center, (ii) A string connecting all 20 images was determined using spline interpolation, (iii) New centers were chosen along the string such that the centers were equidistant along the string. This procedure was repeated until additional iterations produced negligible changes in the restraint centers, indicating string convergence. Plots of image centers over iterations for all string optimizations are shown in Figs. S11-S13. Potentials of mean force (PMFs) were calculated from the final iteration using the multistate Bennett acceptance ratio (MBAR) method as implemented in the pyMBAR package.^67^ Error bars were determined by the standard deviations of PMFs computed using five randomly selected subsets of simulation data from the final iteration.

## Acknowledgments

This research is part of the Blue Waters sustained-petascale computing project, which is supported by the National Science Foundation (Award Nos. OCI-0725070 and ACI-1238993), the State of Illinois, and, as of December 2019, the National Geospatial-Intelligence Agency. Blue Waters is a joint effort of the University of Illinois at Urbana–Champaign and its National Center for Supercomputing Applications. J.C. is a member of the NIH Chemistry-Biology Interface Training Program (T32-GM136629). D.S. acknowledges support from the CAS Fellowship, Center for Advanced Studies at University of Illinois at Urbana–Champaign, and a Sloan Research Fellowship from the Alfred P. Sloan Foundation. D.S. and T.J.D. acknowledge support from the National Institutes of Health Award R35GM142745. This manuscript is the result of funding in whole or in part by the National Institutes of Health (NIH). It is subject to the NIH Public Access Policy. Through acceptance of this federal funding, NIH has been given a right to make this manuscript publicly available in PubMed Central upon the Official Date of Publication, as defined by NIH. The authors thank Austin Weigle for helpful discussion and Matthew Chan for a critical reading of the manuscript.

## Author Contributions

D.S. acquired funding for the project. D.S. and J.C. conceptualized the study. T.J.D. and J.C. performed simulations. T.J.D. and J.C. performed analysis. T.J.D. and J.C. wrote the original draft. D.S. and T.J.D. contributed to manuscript review and editing. D.S. supervised and administered the project.

## Supporting Information

This article includes supporting information: PDF document contains additional details of string optimization protocol and observed mechanisms for all reaction steps.

## Data and Code Availability

In-house code used for simulation setup and data analysis can be found at https://github.com/ShuklaGroup/Strigolactone-Hydrolysis.

