## Supplementary Information for "Mapping Strigolactone Hydrolysis in DWARF14 via QM/MM String Method"

### Contents

**Additional details of string protocol** **S-3**

**Reaction coordinates for each mechanistic step** **S-5**

Table S1: Collective variables harmonically restrained during steered MD and string optimizations. . . . . . . . . . . . . . . . . . . . . . . . . . S-5

**Simulations performed per iteration for each string optimization** **S-6**

Table S2: Simulation time per string iteration for Michael addition pathway S-6

Table S3: Simulation time per string iteration for acyl substitution pathway S-7

**Error bars on potentials of mean force** **S-8**

Table S4: Average PMF values and error bars for Michael addition, initial nucleophilic attack . . . . . . . . . . . . . . . . . . . . . . . . . . S-8

Table S5: Average PMF values and error bars for Michael addition, D-ring separation . . . . . . . . . . . . . . . . . . . . . . . . . . . . . . S-12

Table S6: Average PMF values and error bars for acyl substitution, initial nucleophilic attack . . . . . . . . . . . . . . . . . . . . . . . . . . S-13

Table S7: Average PMF values and error bars for acyl substitution, D-ring separation . . . . . . . . . . . . . . . . . . . . . . . . . . . . . . S-14

Table S8: Average PMF values and error bars for acyl substitution, CLIM formation . . . . . . . . . . . . . . . . . . . . . . . . . . . . . . S-15

Table S9: Average PMF values and error bars for acyl substitution, D-ring-H247 formation from Open D-S97 . . . . . . . . . . . . . . . . . . . . . S-16

Table S10: Average PMF values and error bars for acyl substitution, D-ring–H247 formation from CLIM . . . . . . . . . . . . . . . . . . . . . S-17

**Mechanistic details of non-favored pathways** **S-20**

Michael addition pathway . . . . . . . . . . . . . . . . . . . . . . . . . . . . S-20

Initial nucleophilic attack . . . . . . . . . . . . . . . . . . . . . . . . . . . . . S-20
Figure S1: Distances between key bond forming and breaking atoms during the initial nucleophilic attack in the Michael addition pathway . . . . S-21
D-ring separation . . . . . . . . . . . . . . . . . . . . . . . . . . . . . . . . . S-21
Figure S2: Distances between key bond forming and breaking atoms during the D-ring separation step in the Michael addition pathway . . . . . . S-22
D-ring-H247 formation from CLIM . . . . . . . . . . . . . . . . . . . . . . . . S-22
Figure S3: Distances between key bond forming and breaking atoms during the D-ring-H247 species from CLIM . . . . . . . . . . . . . . . . . . . S-24

**Details of observed reaction mechanisms S-25**

Michael addition pathway . . . . . . . . . . . . . . . . . . . . . . . . . . . . . S-25
Figure S4: Observed mechanism of the initial nucleophilic attack of the Michael addition pathway . . . . . . . . . . . . . . . . . . . . . . . . . . . . . S-25
Figure S5: Observed mechanism of the D-ring separation step of the Michael addition pathway . . . . . . . . . . . . . . . . . . . . . . . . . . . . . S-26
Acyl substitution pathway . . . . . . . . . . . . . . . . . . . . . . . . . . . . . S-27
Figure S6: Observed mechanism of the initial nucleophilic attack of the acyl substitution pathway . . . . . . . . . . . . . . . . . . . . . . . . . . . S-27
Figure S7: Observed mechanism of the D-ring separation step of the acyl substitution pathway . . . . . . . . . . . . . . . . . . . . . . . . . . . S-28
Figure S8: Observed mechanism of the CLIM formation step from the Open D-S97 species of the acyl substitution pathway . . . . . . . . . . . . S-29
Figure S9: Observed mechanism of the D-ring-H247 species from the Open D-S97 species of the acyl substitution pathway . . . . . . . . . . . . S-30
Figure S10: Observed mechanism of the D-ring-H247 species from the CLIM species of the acyl substitution pathway . . . . . . . . . . . . . . . . S-31

**String convergence** **S-32**

Figure S11: Convergence of string restraint centers for acyl substitution (No CLIM path) . . . . . . . . . . . . . . . . . . . . . . . . . . . . . . . S-32

Figure S12: Convergence of string restraint centers for the Michael Addition pathway . . . . . . . . . . . . . . . . . . . . . . . . . . . . . . . . . S-33

Figure S13: Convergence of string restraint centers for acyl substitution (CLIM path) . . . . . . . . . . . . . . . . . . . . . . . . . . . . . . . . . . S-34

#### Additional details of string protocol

To generate an initial pathway for each string, 1 ps steered molecular dynamics trajectories were performed with harmonic restraints on each of the collective variable (CV) sets listed in Table S1 with force constants of 5000 kcal/mol/Å. String optimization was performed using the following procedure:

1. Twenty evenly spaced (in collective variable space) snapshots were extracted from each steered molecular dynamics for each initial string.

2. Each image was simulated for 5 replicates of 100 fs restrained simulations. Harmonic restraints were placed on the same CVs used for steered molecular dynamics simulations with force constants of 100 kcal/mol/Å.

3. Following 100 fs of simulation, the average value of each CV was calculated for each image, and a string in CV space was generated by spline interpolation of average CV values.

4. New restraint centers were chosen by selecting 20 equidistant points along the string. This procedure was repeated for 22-26 iterations. String convergence is shown in Figs. S11-S13.

5. A final refinement iteration was performed to ensure convergence of the string. For this iteration, the number of images was increased such that adjacent restraint centers were spaced by approximately 0.1 Å in CV space. These additional images were generated by arc-length interpolation between the 20 images obtained in the preceding iteration. Harmonic restraint force constants were adjusted on a per-CV basis according to the observed fluctuations in previous iterations: CVs exhibiting minimal positional drift across iterations were restrained with reduced force constants to enhance overlap between neighboring bins, while more flexible CVs retained stronger restraints. This

adaptive restraint scheme was employed to improve statistical overlap and robustness of final MBAR analysis.

### Reaction coordinates for each mechanistic step

Table S1 shows the collective variables to which harmonic restraints were applied for each mechanistic step of both reaction pathways. For all reaction steps, the same collective variables were restrained for both the steered molecular dynamics simulations for initial path generation and string optimization.

Table S1: Collective variables harmonically restrained during steered MD and string optimizations.

| Acyl substitution pathway | | |
|---|---|---|
| Step | Restrained Collective Variables | Physical Process |
| Initial nucleophilic attack | S97:OG-GR24:C16 distance<br>(H247:NE-S97:HG distance) - (S97:OG-S97:HG distance) | S97 attack<br>S97-H247 proton transfer |
| D-ring separation (Open D-S97 formation) | GR24:O3-GR24:C13 distance<br>(GR24:O5-S97:HG distance) - (GR24:O3-S97:HG distance) | D-ring separation<br>Proton transfer |
| CLIM formation | H247:NE-GR24:C13 distance | H247 Nucleophilic attack |
| D-ring-H247 formation from Open D-S97 | H247:NE-GR24:C13 distance<br>GR24:O5-GR24:C16 distance<br>S97:OG-GR24:C16 distance | H247 nucleophilic attack<br>Ring closure<br>Dissociation from S97 |
| D-ring-H247 formation from CLIM | GR24:O5-GR24:C16 distance<br>S97:OG-GR24:C16 distance<br>S97:OG-S97:HG distance | Ring closure<br>Dissociation from S97<br>Proton transfer |
| Michael addition pathway | | |
| Step | Restrained Collective Variables | Physical Process |
| Initial nucleophilic attack | S97:OG-GR24:C12 distance<br>(H247:NE-S97:HG distance) - (S97:OG-S97:HG distance) | S97 attack<br>S97-H247 proton transfer |
| D-ring separation | GR24:O3-GR24:C13 distance<br>GR24:O5-GR24-C16 distance | D-ring separation<br>D-ring opening |

### Simulations performed per iteration for each string optimization

Tables S2-S3 show aggregate simulation time performed for each string iteration for the Michael addition and acyl substitution pathways, respectively. For the CLIM formation step, only a single iteration was performed since string optimization requires restraint of at least two collective variables, and this step only had one restrained collective variable, as indicated in Table S1. Thus, the PMF was computed from a single "iteration" of restrained simulations.

Table S2: Simulation time per string iteration for Michael addition pathway

| Michael addition pathway | | |
|---|---|---|
| Step | Iteration | Simulation time for iteration (ps) |
| Initial nucleophilic attack | 0-3 | 10.0 |
| | 4 | 9.8 |
| | 5-13 | 10.0 |
| | 14 | 8.9 |
| | 15 | 10.0 |
| | 16 | 9.9 |
| | 17-20 | 9.8 |
| | 21-22 | 9.9 |
| | 23 | 40.0 |
| Total (ps) | | 268 |
| D-ring separation | 0-12 | 10.0 |
| | 13 | 5.9 |
| | 14-21 | 4.9 |
| | 22 | 10.5 |
| Total (ps) | | 180 |

Table S3: Simulation time per string iteration for acyl substitution pathway

| Acyl substitution pathway | | |
|---|---|---|
| Step | Iteration | Simulation time for iteration (ps) |
| Initial nucleophilic attack | 0 | 10.0 |
| | 1 | 9.9 |
| | 2 | 10.0 |
| | 3-4 | 9.9 |
| | 5-12 | 10.0 |
| | 13 | 8.0 |
| | 14-23 | 5.7 |
| | 24 | 5.6 |
| | 24-25 | 5.7 |
| | 26 | 20.0 |
| Total (ps) | | 226 |
| D-ring separation | 0-13 | 10.0 |
| | 14 | 7.1 |
| | 15-25 | 5.1 |
| | 26 | 12 |
| Total (ps) | | 215 |
| CLIM formation | 0 | 10.0 |
| | 1 | 46.6 |
| Total (ps) | | 56.6 |
| D-ring-H247 formation from CLIM | 0-4 | 10.0 |
| | 5 | 9.9 |
| | 6-12 | 10.0 |
| | 13 | 9.8 |
| | 14-26 | 10.0 |
| | 27 | 150.0 |
| Total (ps) | | 420 |
| D-ring-H247 formation from Open D-S97 | 0-4 | 10.0 |
| | 5 | 9.2 |
| | 6 | 8.3 |
| | 7-8 | 10.0 |
| | 10 | 9.9 |
| | 11 | 9.8 |
| | 12 | 9.9 |
| | 13 | 9.8 |
| | 14-26 | 10.0 |
| | 27 | 100 |
| Total (ps) | | 367 |

### Error bars on potentials of mean force

Tables S4-S10 show the average and standard deviations of PMF values across five randomly subsampled sets of data for each mechanistic step. Each subsampled data set contained 80% of the data from the final string iteration.

Table S4: Average PMF values and error bars for Michael addition, initial nucleophilic attack

| Michael addition, initial nucleophilic attack | | |
|---|---|---|
| Image | Average free energy (kcal/mol) | Error (kcal/mol) |
| 0 | 0.00 | 0.00 |
| 1 | -0.25 | 0.00 |
| 2 | -0.36 | 0.00 |
| 3 | -0.38 | 0.00 |
| 4 | -0.36 | 0.00 |
| 5 | -0.36 | 0.00 |
| 6 | -0.35 | 0.00 |
| 7 | -0.31 | 0.01 |
| 8 | -0.21 | 0.01 |
| 9 | -0.08 | 0.01 |
| 10 | 0.05 | 0.01 |
| 11 | 0.18 | 0.01 |
| 12 | 0.33 | 0.01 |
| 13 | 0.49 | 0.01 |
| 14 | 0.66 | 0.01 |
| 15 | 0.81 | 0.01 |
| 16 | 0.91 | 0.01 |
| 17 | 0.97 | 0.01 |
| 18 | 1.00 | 0.01 |

| Image | Average free energy (kcal/mol) | Error (kcal/mol) |
|---|---|---|
| 19 | 0.98 | 0.01 |
| 20 | 0.93 | 0.01 |
| 21 | 0.86 | 0.01 |
| 22 | 0.71 | 0.01 |
| 23 | 0.61 | 0.01 |
| 24 | 0.59 | 0.01 |
| 25 | 0.66 | 0.01 |
| 26 | 0.82 | 0.01 |
| 27 | 1.02 | 0.02 |
| 28 | 1.22 | 0.02 |
| 29 | 1.46 | 0.02 |
| 30 | 1.84 | 0.02 |
| 31 | 2.35 | 0.02 |
| 32 | 2.96 | 0.02 |
| 33 | 3.66 | 0.02 |
| 34 | 4.45 | 0.02 |
| 35 | 5.27 | 0.02 |
| 36 | 6.05 | 0.02 |
| 37 | 6.75 | 0.02 |
| 38 | 7.35 | 0.02 |
| 39 | 7.90 | 0.03 |
| 40 | 8.45 | 0.03 |
| 41 | 9.01 | 0.03 |
| 42 | 9.54 | 0.03 |
| 43 | 10.05 | 0.03 |
| 44 | 10.60 | 0.03 |

| Image | Average free energy (kcal/mol) | Error (kcal/mol) |
|---|---|---|
| 45 | 11.18 | 0.03 |
| 46 | 11.72 | 0.03 |
| 47 | 12.14 | 0.03 |
| 48 | 12.44 | 0.03 |
| 49 | 12.68 | 0.03 |
| 50 | 12.88 | 0.03 |
| 51 | 13.04 | 0.03 |
| 52 | 13.24 | 0.03 |
| 53 | 13.51 | 0.03 |
| 54 | 13.82 | 0.03 |
| 55 | 13.95 | 0.04 |
| 56 | 13.96 | 0.03 |
| 57 | 13.97 | 0.03 |
| 58 | 13.92 | 0.04 |
| 59 | 13.84 | 0.04 |
| 60 | 13.84 | 0.04 |
| 61 | 13.91 | 0.04 |
| 62 | 14.04 | 0.04 |
| 63 | 14.23 | 0.04 |
| 64 | 14.44 | 0.04 |
| 65 | 14.63 | 0.03 |
| 66 | 14.80 | 0.03 |
| 67 | 14.98 | 0.03 |
| 68 | 15.19 | 0.03 |
| 69 | 15.48 | 0.04 |
| 70 | 15.81 | 0.04 |

| Image | Average free energy (kcal/mol) | Error (kcal/mol) |
|---|---|---|
| 71 | 15.93 | 0.04 |
| 72 | 15.67 | 0.04 |
| 73 | 15.21 | 0.04 |
| 74 | 14.57 | 0.04 |
| 75 | 13.68 | 0.04 |
| 76 | 12.74 | 0.04 |
| 77 | 11.97 | 0.04 |
| 78 | 11.49 | 0.04 |
| 79 | 11.30 | 0.04 |

Table S5: Average PMF values and error bars for Michael addition, D-ring separation

| Michael addition, D-ring separation | | |
|---|---|---|
| Image | Average free energy (kcal/mol) | Error (kcal/mol) |
| 0 | 0.00 | 0.00 |
| 1 | 0.74 | 0.01 |
| 2 | 2.97 | 0.01 |
| 3 | 5.40 | 0.03 |
| 3.5 | 5.63 | 0.03 |
| 4 | 5.48 | 0.03 |
| 5 | 3.21 | 0.03 |
| 6 | 1.16 | 0.03 |
| 7 | -0.14 | 0.02 |
| 8 | -1.09 | 0.03 |
| 9 | -1.93 | 0.03 |
| 10 | -2.16 | 0.03 |
| 11 | -2.00 | 0.03 |
| 12 | -1.22 | 0.02 |
| 13 | -0.92 | 0.03 |
| 14 | -0.88 | 0.04 |
| 15 | -0.90 | 0.04 |
| 16 | -0.90 | 0.05 |
| 17 | -0.95 | 0.05 |
| 18 | -0.99 | 0.06 |
| 19 | -0.92 | 0.07 |

Table S6: Average PMF values and error bars for acyl substitution, initial nucleophilic attack

| Acyl substitution, initial nucleophilic attack | | |
|---|---|---|
| Image | Average free energy (kcal/mol) | Error (kcal/mol) |
| 0 | 0.00 | 0.00 |
| 1 | -0.24 | 0.00 |
| 2 | -0.18 | 0.00 |
| 3 | 0.10 | 0.00 |
| 4 | 0.46 | 0.01 |
| 5 | 0.81 | 0.01 |
| 6 | 0.98 | 0.01 |
| 7 | 1.01 | 0.01 |
| 8 | 1.06 | 0.01 |
| 9 | 1.20 | 0.01 |
| 10 | 1.40 | 0.01 |
| 11 | 1.64 | 0.01 |
| 12 | 1.78 | 0.01 |
| 13 | 1.87 | 0.02 |
| 14 | 2.10 | 0.02 |
| 15 | 2.46 | 0.02 |
| 16 | 2.77 | 0.02 |
| 17 | 3.13 | 0.02 |
| 18 | 3.62 | 0.02 |
| 19 | 4.28 | 0.02 |
| 20 | 5.01 | 0.03 |
| 21 | 5.92 | 0.03 |
| 22 | 6.80 | 0.03 |
| 23 | 7.19 | 0.03 |
| 24 | 7.72 | 0.02 |
| 25 | 8.28 | 0.02 |
| 26 | 8.66 | 0.02 |
| 27 | 9.06 | 0.02 |
| 28 | 9.74 | 0.02 |
| 29 | 10.69 | 0.02 |
| 30 | 11.69 | 0.02 |
| 31 | 12.74 | 0.02 |
| 32 | 13.49 | 0.02 |
| 33 | 13.95 | 0.02 |
| 34 | 13.69 | 0.02 |
| 35 | 13.00 | 0.02 |
| 36 | 12.68 | 0.02 |
| 37 | 12.60 | 0.02 |
| 38 | 12.62 | 0.02 |
| 39 | 12.69 | 0.02 |

Table S7: Average PMF values and error bars for acyl substitution, D-ring separation

| Acyl substitution, D-ring separation (Open D-S97 formation) | | |
|---|---|---|
| Image | Average free energy (kcal/mol) | Error (kcal/mol) |
| 0 | 0.00 | 0.00 |
| 1 | 0.14 | 0.00 |
| 2 | 0.95 | 0.01 |
| 3 | 1.96 | 0.00 |
| 4 | 3.30 | 0.00 |
| 5 | 5.42 | 0.00 |
| 5.5 | 6.78 | 0.00 |
| 6 | 8.25 | 0.00 |
| 6.5 | 9.25 | 0.00 |
| 7 | 9.68 | 0.00 |
| 7.5 | 9.81 | 0.01 |
| 8 | 9.52 | 0.01 |
| 9 | 8.35 | 0.01 |
| 9.5 | 7.66 | 0.01 |
| 10 | 6.98 | 0.01 |
| 11 | 5.71 | 0.01 |
| 12 | 4.69 | 0.01 |
| 13 | 3.68 | 0.01 |
| 14 | 3.13 | 0.01 |
| 15 | 2.97 | 0.01 |
| 16 | 3.00 | 0.02 |
| 17 | 3.04 | 0.02 |
| 18 | 3.05 | 0.02 |
| 19 | 3.12 | 0.02 |

Table S8: Average PMF values and error bars for acyl substitution, CLIM formation

| Acyl substitution, CLIM formation | | |
|---|---|---|
| Image | Average free energy (kcal/mol) | Error (kcal/mol) |
| 0 | 0.00 | 0.00 |
| 1 | -0.17 | 0.00 |
| 2 | -0.20 | 0.00 |
| 3 | -0.17 | 0.00 |
| 4 | -0.11 | 0.00 |
| 5 | 0.00 | 0.00 |
| 6 | 0.17 | 0.00 |
| 7 | 0.48 | 0.00 |
| 8 | 0.89 | 0.00 |
| 9 | 1.23 | 0.01 |
| 10 | 1.79 | 0.01 |
| 11 | 2.46 | 0.01 |
| 12 | 3.10 | 0.01 |
| 13 | 3.77 | 0.01 |
| 14 | 4.89 | 0.01 |
| 15 | 5.80 | 0.01 |
| 16 | 6.15 | 0.01 |
| 17 | 5.77 | 0.01 |
| 18 | 5.61 | 0.01 |
| 19 | 5.71 | 0.01 |

Table S9: Average PMF values and error bars for acyl substitution, D-ring-H247 formation from Open D-S97

| Acyl substitution, D-ring-H247 formation from Open D-S97 | | |
|---|---|---|
| Image | Average free energy (kcal/mol) | Error (kcal/mol) |
| 0 | 0.00 | 0.00 |
| 1 | -0.04 | 0.00 |
| 2 | 0.07 | 0.00 |
| 3 | 0.29 | 0.00 |
| 4 | 0.58 | 0.00 |
| 5 | 0.90 | 0.00 |
| 6 | 1.22 | 0.00 |
| 7 | 1.50 | 0.00 |
| 8 | 1.72 | 0.00 |
| 9 | 1.93 | 0.00 |
| 10 | 2.17 | 0.00 |
| 11 | 2.42 | 0.00 |
| 12 | 2.65 | 0.00 |
| 13 | 2.87 | 0.00 |
| 14 | 3.10 | 0.00 |
| 15 | 3.37 | 0.00 |
| 16 | 3.69 | 0.00 |
| 17 | 4.05 | 0.00 |
| 18 | 4.44 | 0.00 |
| 19 | 4.85 | 0.01 |
| 20 | 5.26 | 0.01 |
| 21 | 5.70 | 0.01 |
| 22 | 6.18 | 0.01 |
| 23 | 6.68 | 0.01 |
| 24 | 7.19 | 0.01 |
| 25 | 7.62 | 0.01 |
| 26 | 7.89 | 0.01 |
| 27 | 7.99 | 0.00 |
| 28 | 8.03 | 0.00 |
| 29 | 8.05 | 0.01 |
| 30 | 8.11 | 0.01 |
| 31 | 8.25 | 0.01 |
| 32 | 8.53 | 0.01 |
| 33 | 8.95 | 0.01 |
| 34 | 9.57 | 0.01 |
| 35 | 10.22 | 0.01 |
| 36 | 10.81 | 0.01 |
| 37 | 11.22 | 0.01 |
| 38 | 11.39 | 0.01 |
| 39 | 11.46 | 0.01 |

Table S10: Average PMF values and error bars for acyl substitution, D-ring–H247 formation from CLIM

| Acyl substitution, D-ring–H247 formation from CLIM | | |
|---|---|---|
| Image | Average free energy (kcal/mol) | Error (kcal/mol) |
| 0 | 0.00 | 0.00 |
| 1 | -0.02 | 0.00 |
| 2 | 0.09 | 0.00 |
| 3 | 0.32 | 0.00 |
| 4 | 0.64 | 0.00 |
| 5 | 1.01 | 0.00 |
| 6 | 1.39 | 0.00 |
| 7 | 1.77 | 0.00 |
| 8 | 2.18 | 0.00 |
| 9 | 2.65 | 0.00 |
| 10 | 3.20 | 0.00 |
| 11 | 3.79 | 0.00 |
| 12 | 4.38 | 0.00 |
| 13 | 4.99 | 0.00 |
| 14 | 5.63 | 0.00 |
| 15 | 6.31 | 0.00 |
| 16 | 7.05 | 0.01 |
| 17 | 7.82 | 0.01 |
| 18 | 8.65 | 0.01 |
| 19 | 9.53 | 0.01 |
| 20 | 10.49 | 0.00 |
| 21 | 11.53 | 0.00 |
| 22 | 12.63 | 0.00 |

| Image | Average free energy (kcal/mol) | Error (kcal/mol) |
|---|---|---|
| 23 | 13.73 | 0.00 |
| 24 | 14.79 | 0.00 |
| 25 | 15.81 | 0.00 |
| 26 | 16.74 | 0.00 |
| 27 | 17.61 | 0.00 |
| 28 | 18.39 | 0.00 |
| 29 | 19.08 | 0.00 |
| 30 | 19.79 | 0.00 |
| 31 | 20.53 | 0.00 |
| 32 | 21.23 | 0.00 |
| 33 | 21.80 | 0.00 |
| 34 | 22.23 | 0.00 |
| 35 | 22.40 | 0.00 |
| 36 | 22.47 | 0.00 |
| 37 | 22.46 | 0.00 |
| 38 | 22.40 | 0.00 |
| 39 | 22.30 | 0.00 |
| 40 | 22.16 | 0.01 |
| 41 | 22.02 | 0.01 |
| 42 | 21.91 | 0.00 |
| 43 | 21.84 | 0.00 |
| 44 | 21.83 | 0.00 |
| 45 | 21.88 | 0.00 |
| 46 | 22.02 | 0.00 |
| 47 | 22.24 | 0.01 |
| 48 | 22.56 | 0.01 |

| Image | Average free energy (kcal/mol) | Error (kcal/mol) |
|---|---|---|
| 49 | 22.96 | 0.01 |
| 50 | 23.45 | 0.01 |
| 51 | 24.04 | 0.01 |
| 52 | 24.74 | 0.01 |
| 53 | 25.48 | 0.01 |
| 54 | 25.48 | 0.01 |
| 55 | 24.80 | 0.01 |
| 56 | 24.21 | 0.01 |
| 57 | 23.82 | 0.01 |
| 58 | 23.62 | 0.01 |
| 59 | 23.61 | 0.01 |

### Mechanistic details of non-favored pathways

#### Michael addition pathway

##### Initial nucleophilic attack

The Michael addition pathway begins with a proton transfer-coupled nucleophilic attack of S97 on the enol-ether bridge (Fig. S1). In contrast to the initial nucleophilic attack in the acyl substitution pathway, the optimized reaction pathway is more stepwise than concerted, indicated by the protonation of H247 well before the formation of the S97-GR24 bond. Arrow-pushing details of this step are shown in Fig. S4.

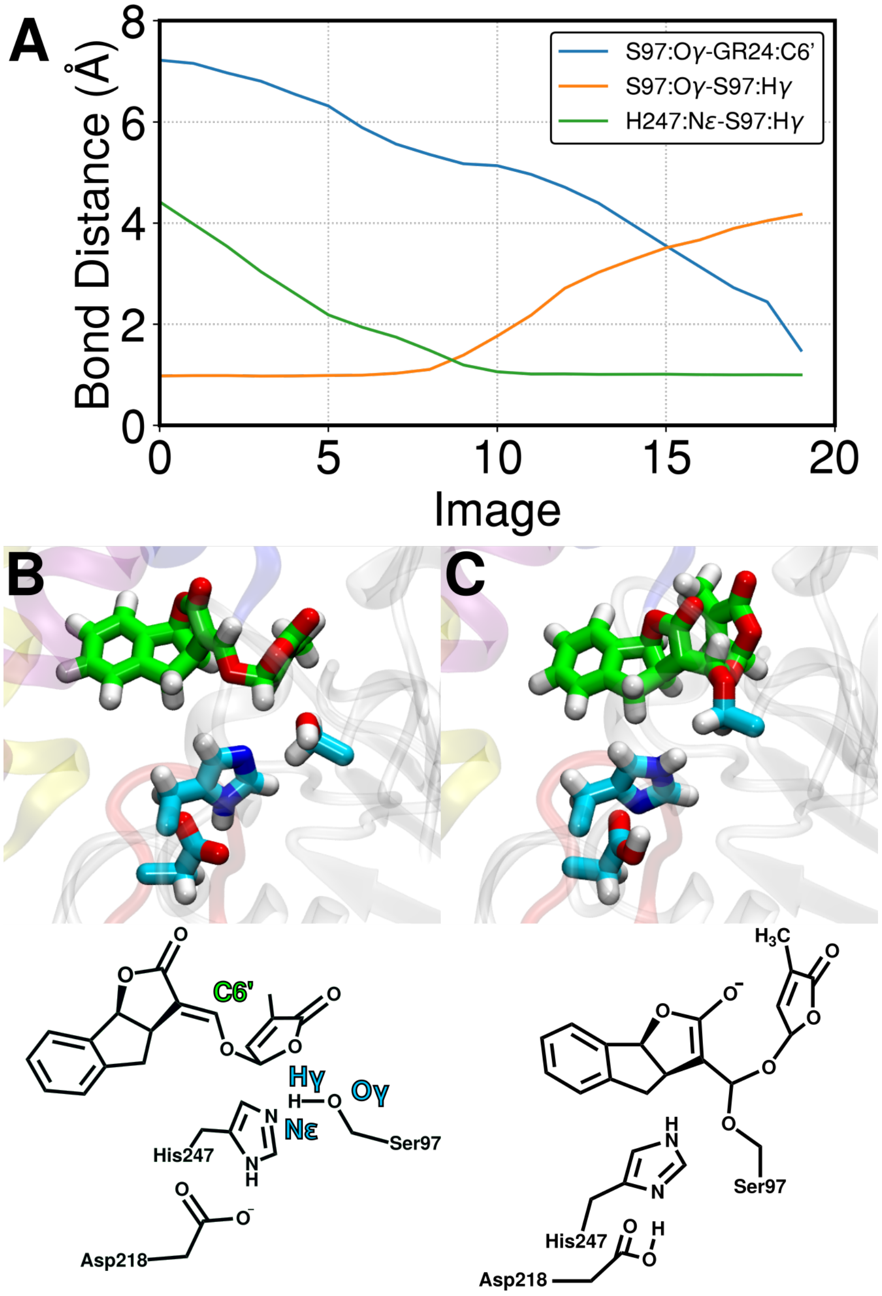


Figure S1: Distances between key bond forming and breaking atoms during the initial nucleophilic attack in the Michael addition pathway (A). The structures of (B) the initial state (intact GR24 and the catalytic triad) and (C) the final state (tetrahedral intermediate with S97 bound to enol-ether bridge) are shown. Since the PMF profile for this step is monotonically increasing, no transition state is shown.

**D-ring separation**

The D-ring separation step of the Michael addition pathway occurs in conjunction with opening of the D-ring. Arrow-pushing details of this step are shown in Fig. S5.

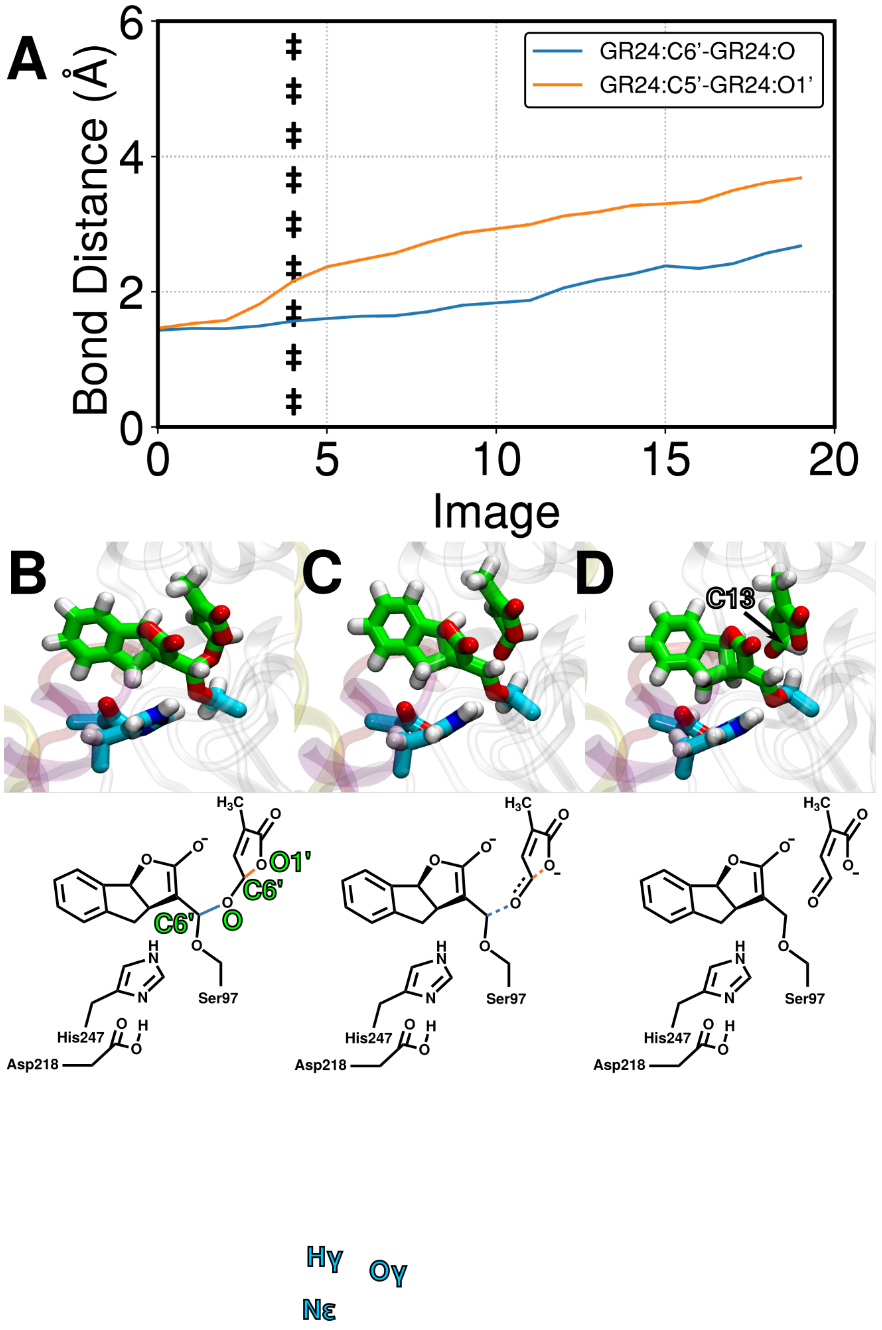


Figure S2: Distances between key bond forming and breaking atoms during the D-ring separation step in the Michael addition pathway (A). The structures of (B) the initial state (tetrahedral intermediate from previous step), (C) transition state, defined as the maximum point of the PMF, and (D) the final state (ABC-scaffold bound to S97, separated D-ring) are shown.

#### D-ring-H247 formation from CLIM

In addition to forming directly from the Open D-S97 intermediate, the D-ring-H247 species can form from a CLIM species, albeit via a higher energy pathway. Key bonds that form and break and key intermediate and transitional species are shown in Fig. S3. Arrow-pushing details of this reaction step are shown in Fig. S10.

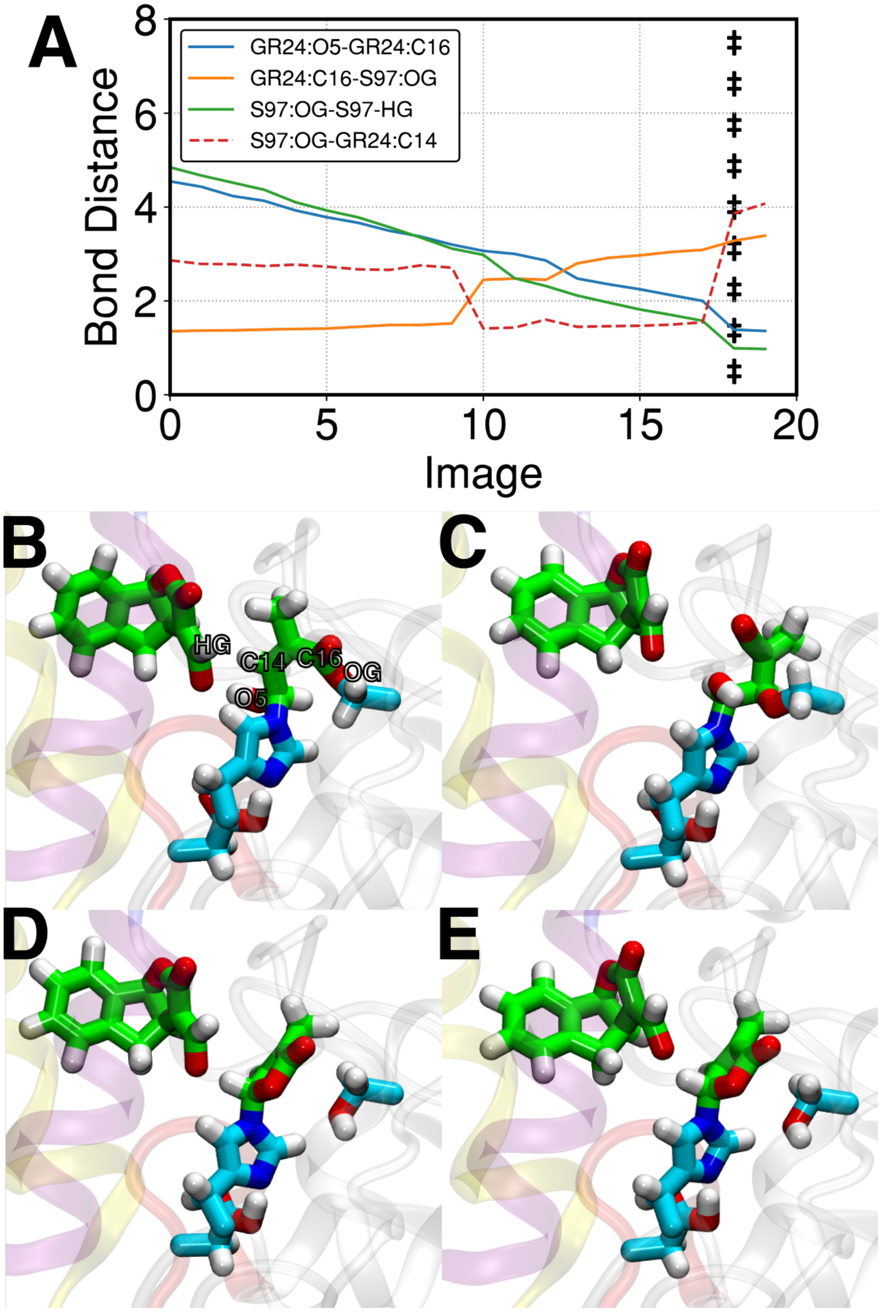


Figure S3: Distances between key bond forming and breaking atoms during the D-ring-H247 species from CLIM (A). Structures shown are (B) the initial CLIM state, (C) an intermediate species along the pathway in which S97 is bound to the C14 carbon on the D-ring which forms at image 10 of the string, (D) the maximum point on the PMF profile for this step, and (E) the final D-ring-H247 modification.

### Details of observed reaction mechanisms

#### Michael addition pathway

Figs. S4-S5 show arrow-pushing schemes for observed Michael addition pathway with corresponding snapshots from simulations.

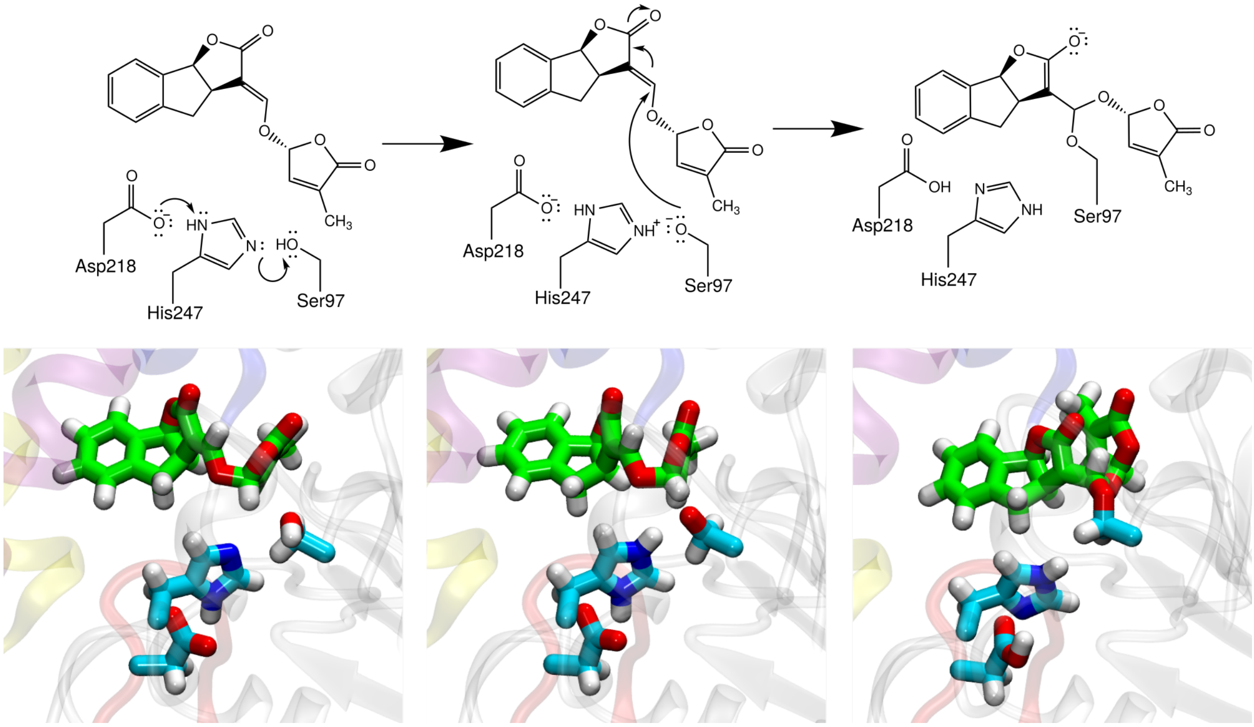


Figure S4: Observed mechanism of the initial nucleophilic attack of the Michael addition pathway. Corresponding snapshots of simulations are shown below each 2-dimensional schematic. This observed mechanism differs from the one shown in Main Text, Fig. 2A in that the proton transfer and nucleophilic attack occur in a stepwise manner.

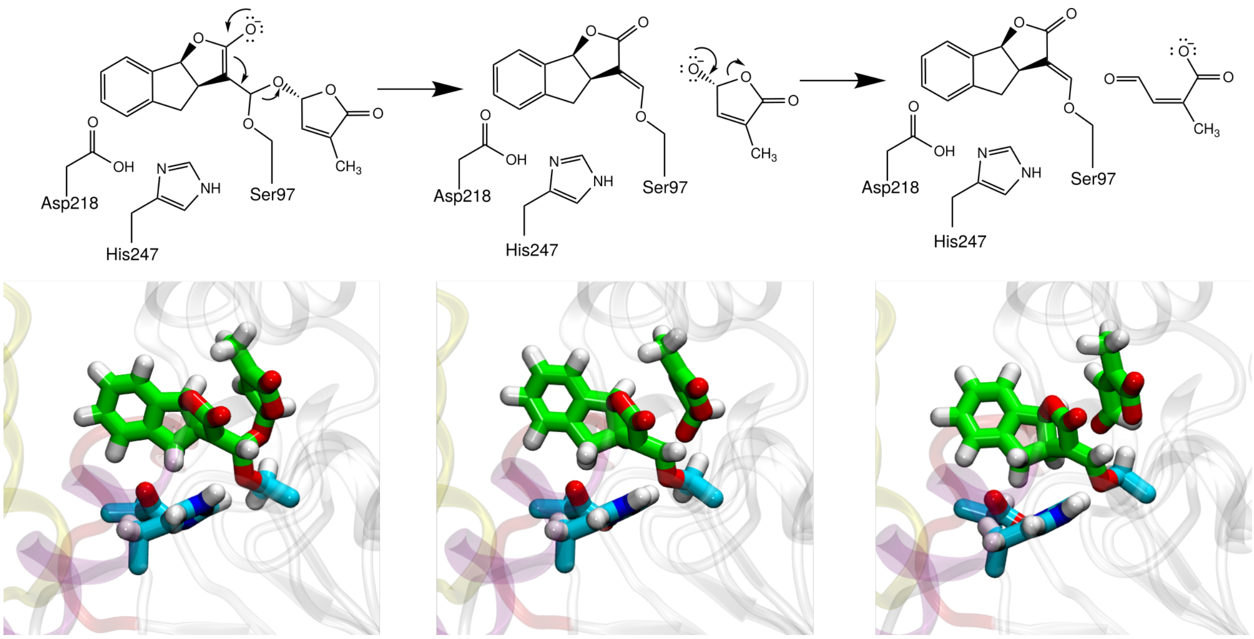


Figure S5: Observed mechanism of the D-ring separation step of the Michael addition pathway. Corresponding snapshots of simulations are shown below each 2-dimensional schematic. This observed mechanism differs from the one shown in Main Text, Fig. 2A in that the separation of the D-ring is coupled with opening of the D-ring.

### Acyl substitution pathway

Figs. S6-S10 show arrow-pushing schemes for observed acyl substitution pathway with corresponding snapshots from simulations.

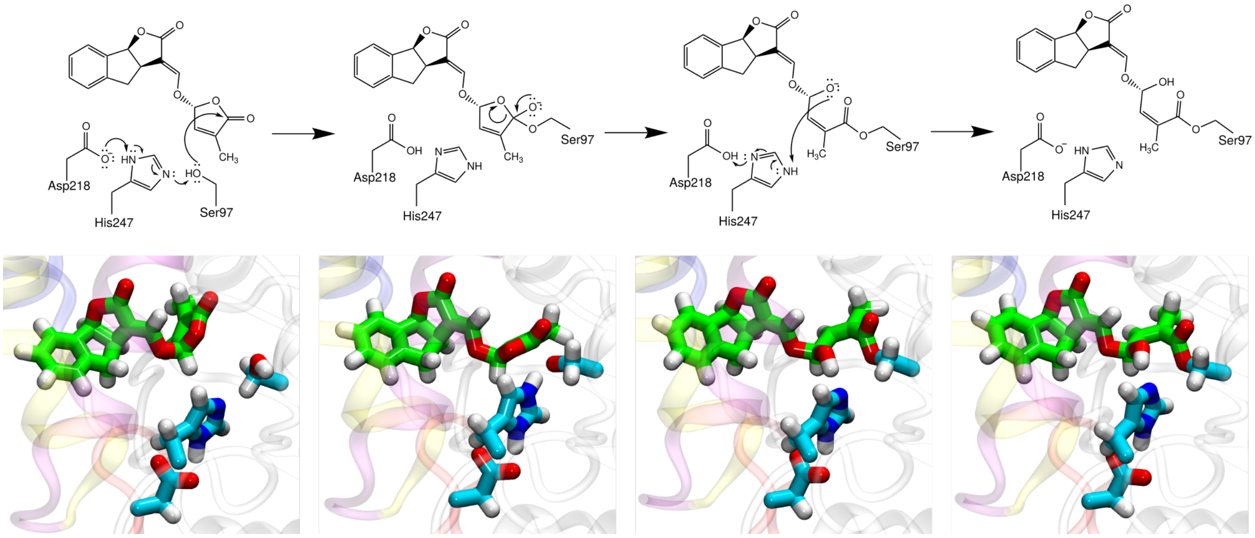


Figure S6: Observed mechanism of the initial nucleophilic attack of the acyl substitution pathway. Corresponding snapshots of simulations are shown below each 2-dimensional schematic. This observed mechanism differs from the one shown in Main Text, Fig. 2B in that nucleophilic attack of the D-ring results in a spontaneos ring-opening.

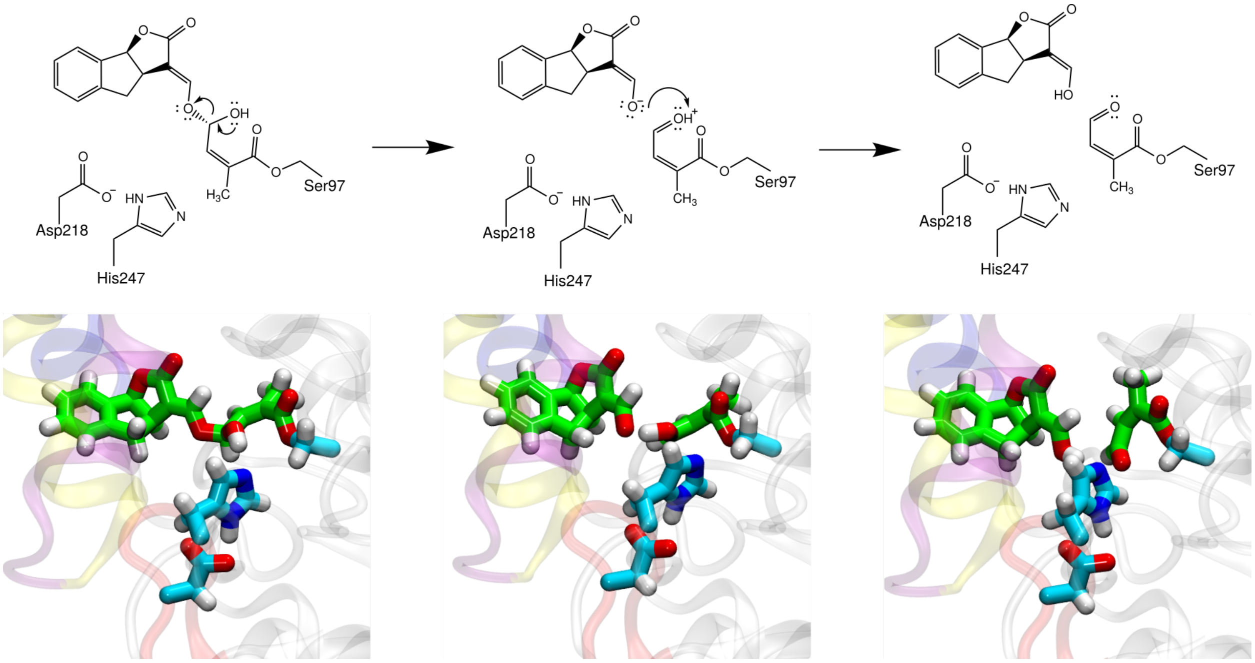


Figure S7: Observed mechanism of the D-ring separation step of the acyl substitution pathway. Corresponding snapshots of simulations are shown below each 2-dimensional schematic. This observed mechanism differs from the one shown in Main Text, Fig. 2B in that the D-ring is opened prior to detachment, however, the species formed by this step (Open D-S97) matches that which is formed by the D-ring detachment step shown in Fig. 2B.

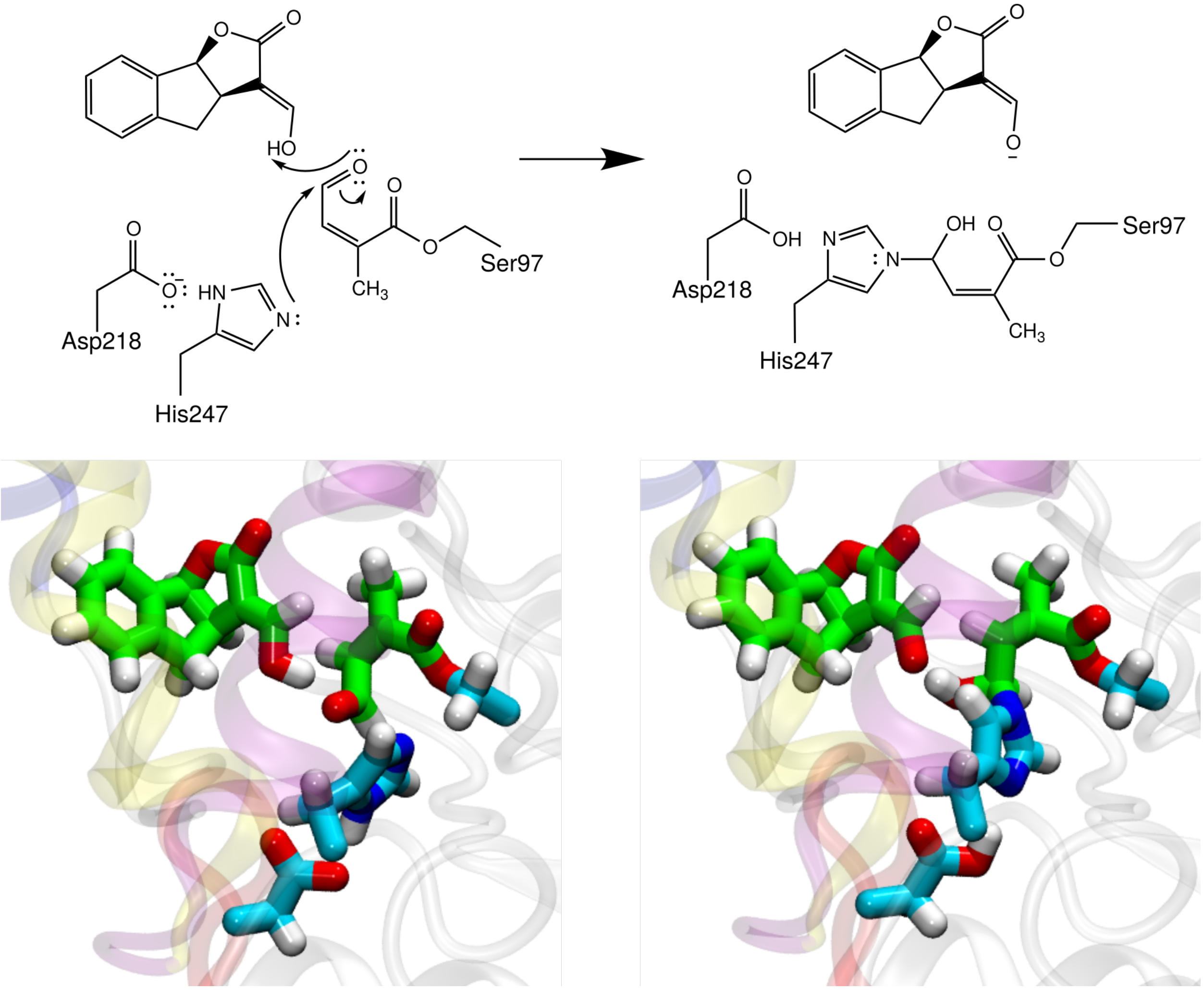


Figure S8: Observed mechanism of the CLIM formation step from the Open D-S97 species of the acyl substitution pathway. Corresponding snapshots of simulations are shown below each 2-dimensional schematic. This observed mechanism matches the one shown in Main Text, Fig. 2B.

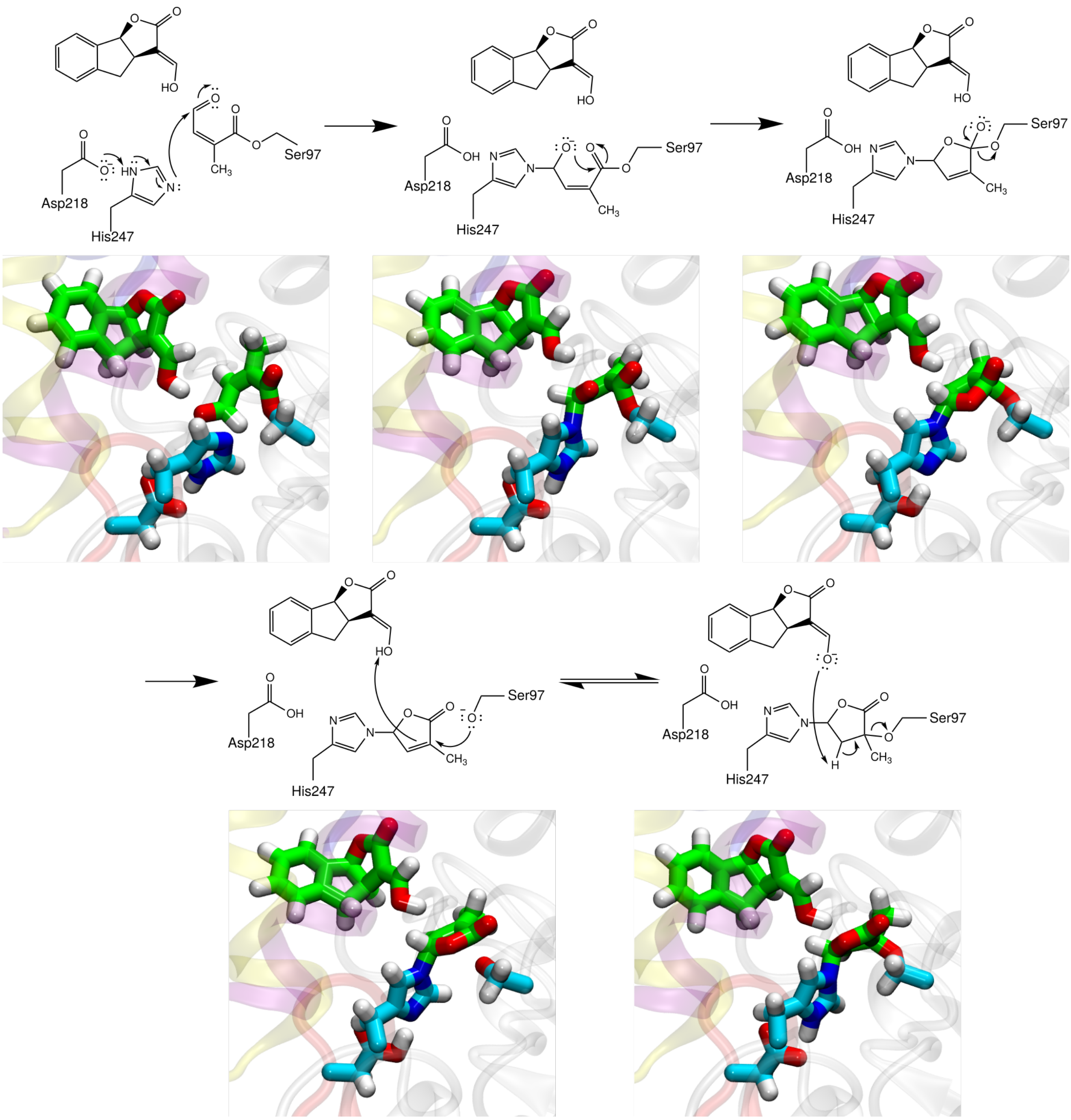


Figure S9: Observed mechanism of the D-ring-H247 species from the Open D-S97 species of the acyl substitution pathway. Corresponding snapshots of simulations are shown below each 2-dimensional schematic. This observed mechanism differs from the one shown in Main Text, Fig. 2B in that the D-ring-H247 species is formed in a concerted manner rather than forming a stable CLIM species. While a transient CLIM-like species is formed (second panel), it differs from the stable CLIM species in the orientation and protonation state of the GR24:O5 oxygen.

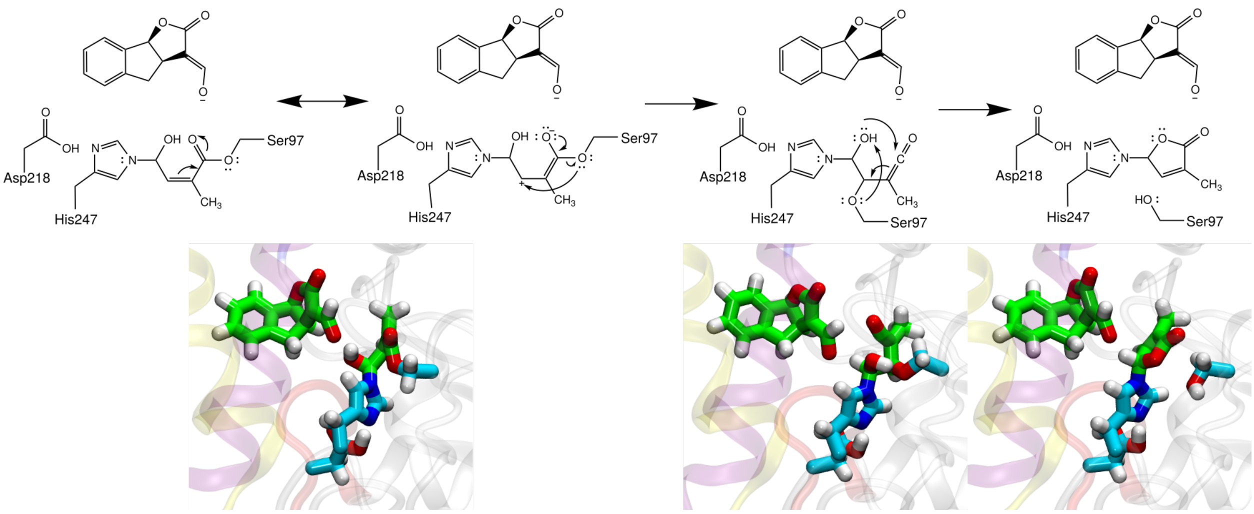


Figure S10: Observed mechanism of the D-ring-H247 species from the CLIM species of the acyl substitution pathway. Corresponding snapshots of simulations are shown below each 2-dimensional schematic. This observed mechanism differs from the one shown in Main Text, Fig. 2B in that rather than the GR24:O5 oxygen directly attacking the GR24:C16 carbon while the D-ring is bound to both S97 and H247, the D-ring dissociations from S97 and reassociates with S97 via a different carbon atom on the ring, facilitating the ring closure reaction.

### String convergence

Figs. S11-S13 show the convergence of restraint centers along strings for each string optimization. Strings images are shown projected onto each pair of collective variables constrained during the respective string optimization.

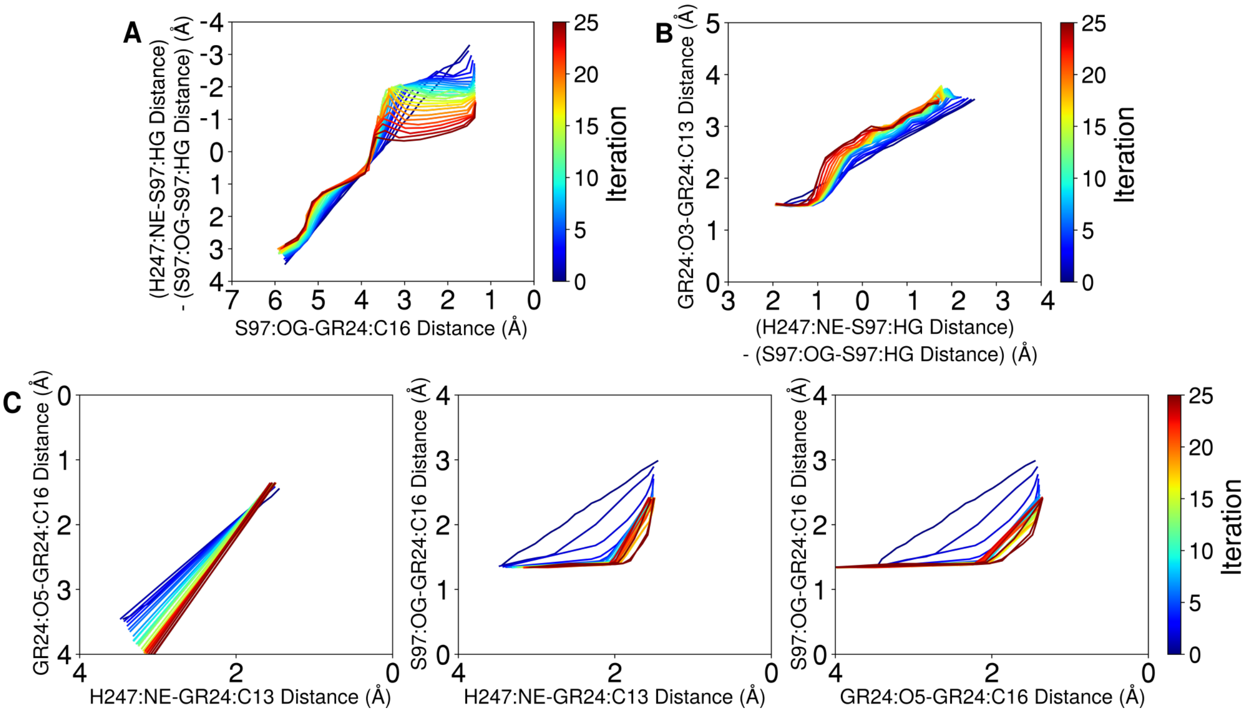


Figure S11: Convergence of string restraint centers for (A) the initial nucleophilic attack step, (B) the D-ring separation step, and (C) the formation of the D-ring-H247 from Open-D-S97 species of the acyl substitution reaction pathway projected onto restrained collective variables for each respective step. Blue lines show restraint centers along the string for earlier iterations, and red lines show later iterations.

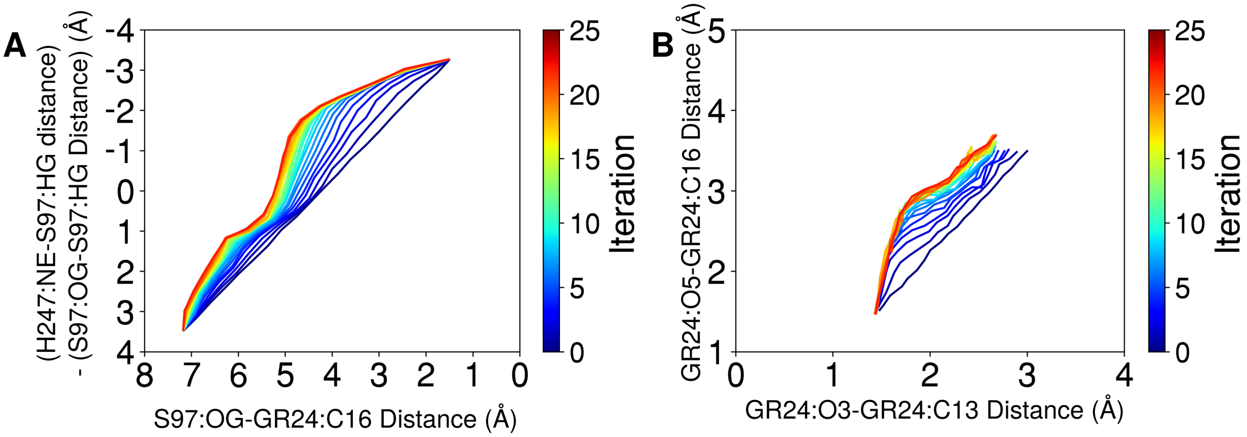


Figure S12: Convergence of string restraint centers for (A) the initial nucleophilic attack step and (B) the D-ring separation step of the Michael addition reaction pathway projected onto restrained collective variables for each respective step. Blue lines show restraint centers along the string for earlier iterations, and red lines show later iterations.

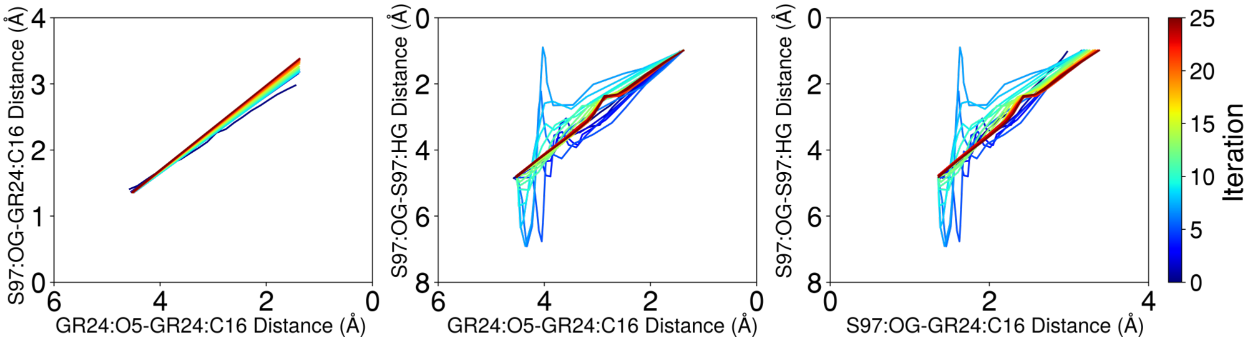


Figure S13: Convergence of string restraint centers for the formation of the D-ring-H247 from the CLIM species of the acyl substitution reaction pathway projected onto each pair of restrained collective variables. Blue lines show restraint centers along the string for earlier iterations, and red lines show later iterations.
